# MicroRNA397 regulates tolerance to drought and fungal infection by regulating lignin deposition in chickpea root

**DOI:** 10.1101/2022.12.12.520062

**Authors:** Nilesh Kumar Sharma, Santosh Kumar Gupta, Vadivelmurugan Irulappan, Shalini Yadav, Aleena Francis, Muthappa Senthil-Kumar, Debasis Chattopadhyay

## Abstract

Plants deposit lignin in the secondary cell wall as a common response to drought and pathogen attacks. Cell wall localized multicopper oxidase family enzymes LACCASES (LACs) catalyze the formation of monolignol radicals and facilitate lignin formation. We show an upregulation of the expression of several *LAC* genes and a downregulation of microRNA397 (CamiR397) in response to natural drought in chickpea roots. CamiR397 was found to target *LAC4* and *LAC17L* out of twenty annotated *LAC*s in chickpea. CamiR397 and its target genes are expressed in root. Overexpression of CamiR397 reduced expression of *LAC4* and *LAC17L* and lignin deposition in chickpea root xylem causing reduction in xylem diameter. Downregulation of CamiR397 activity by expressing a short tandem target mimic (STTM397) construct increased root lignin deposition in chickpea. CamiR397-overexpressing (miR397OX) and STTM397 chickpea lines showed sensitivity and tolerance, respectively, to drought. Infection with a fungal pathogen *Macrophomina phaseolina*, responsible for dry root rot disease in chickpea, induced local lignin deposition and *LAC* gene expression. CamiR397-overexpressing and STTM397 chickpea lines showed more sensitivity and tolerance, respectively, to dry root rot. Our results demonstrated the regulatory role of CamiR397 in root lignification during drought and dry root rot in an agriculturally important crop chickpea.

## Introduction

Being sessile, plants, especially the land plants must endure the environmental stresses to survive. The plant adapts to low soil moisture condition by deeper penetration of roots the in soil in search of moisture and facilitating efficient water movement in the main root (Fischer *et al*., 1982). Abiotic stress adaptation includes shift in allometry of the plants through maintaining stomatal conductance for less water consumption and, deeper and diverse root profile into the soil (Mencuccini, 2003; Addington *et al*., 2006; Maseda and Fernandez, 2006). The drop in lateral growth is very usual under harsh environmental conditions as adaptation by the plants (Xiong *et al*., 2006). The water and mineral transport become crucial under water scarce condition. Plants have well developed vascular tissue systems *viz*. xylem and phloem. Water and minerals are transported by xylem, whilst soluble organic compounds (photosynthate) are carried via phloem (Lalonde *et al*., 2004).

Xylem vessel-based passive system of water transport is particularly crucial for the survival of land plants. Besides the transport, xylem provides mechanical strength to the root for physical support to the plants (Turner *et al*., 2007; Gibson, 2012). Many biomolecules travel through the tracheary tissues (Tracheary Elements, TEs) to several destinations in the plant (Kim *et al*., 2014). Most of the xylem tissue is non-living and does not have any capacity to adapt to changes in hydraulic demand (Timothy JB, 2009). Xylem vessels are lignified which provides strength and impermeable cell wall for leakproof water and mineral transport (Fukuda, 2000).

Lignification is a product of oxidative polymerization of different types of aromatic monomers biosynthesized from phenylalanine, known as monolignols *viz*. p-coumaryl alcohol, coniferyl alcohol and sinapyl alcohol. These monomers produce three main lignin subunits: H (p-hydroxylphenyl), G (guaiacyl) and S (syringyl), respectively (Boerjan *et al*., 2003; Ralph *et al*., 2004), which get integrated into the polymer and coupled with the polysaccharide chain in the cell wall. These different monomers govern the recurrence of multiple chemical bonds in the polymer. There was no significant impact on plant development when the level of G and S lignin units was changed. These compositional alterations in the types of monomer units result in structural changes in the polymerization of lignin and, therefore, on cell wall properties (Vanholme *et al*., 2010). Lignin deposition is not equal in all the plant parts, and it is rather a regulated process. Drought in plants lead to enhanced lignin deposition in the xylem vessels of plants (Caño-Delgado *et al*., 2003; Tronchet *et al*., 2010; Liu *et al*., 2018). *Leucaena leucocephala* exhibited increased lignin deposition in the developing root under drought and salinity stress (Srivastava *et al*., 2015). In soyabean, lignin content was increased in the root elongation zone under drought stress, and this increased lignin deposition caused a reduction in the elongation of cell wall in the elongation zone (Yamaguchi *et al*., 2010).

The study available on plant lignification is very much limited to the enzymes of lignin biosynthesis pathways. The PEROXIDASE group of enzymes garnered a lot of attention in the research on lignification. Another group of enzymes involved in the polymerization of lignol monomers is the LACCASES. LACCASES (LACs) are a group of multi-copper glycoprotein oxidase (*p*-diphenol oxidoreductase or urishiol oxidase EC 1.10.3.2) which can catalyze several phenolic, inorganic and/or aromatic amine substrates. Unlike PEROXIDASE, these oxygen dependent enzymes do not need H_2_O_2_ which has short life span making it a rate limiting factor (Hüttermann *et al*., 2001; Tobimatsu and Schuetz, 2019). LACCASES are present in bacteria, fungi, higher plants (Givaudan *et al*., 1993; Diamantidis *et al*., 2000), and in arthropods (Thomas *et al*., 1989; Cardenas and Dankert, 2000). Most studied LACCASES are fungal LACCASES which catalyse degradation of lignin (Thurston, 1994). LACCASES in fungi acts through hydrolyzing lignin polymers in parallel to PEROXIDASE enzyme activity (Eggert *et al*., 1997; Zhao *et al*., 2013). In *Arabidopsis thaliana*, there are a total of 17 LAC-encoding genes reported with a few genes showing distinctive expression patterns (Cai *et al*., 2006; Turlapati *et al*., 2011).

*AtLAC4* expresses specifically in the vascular bundles, interfascicular fibers and seed coat columella while *AtLAC7* in the hydathodes and root hairs, *AtLAC8* in pollen grains and phloem, *AtLAC15* in seed coat cell walls, and *AtLAC17* in interfascicular fibers (Turlapati *et al*., 2011; Berthert *et al*., 2011). In *Saccharum officinarum, SofLAC* showed differential expression in sclerenchyma bundles and parenchyma cells adjoining the vascular bundles of young internodes (Cesarino *et al*., 2013). Single or double mutant of *lac4* and *lac17* lead to a minor and major reduction in lignification, respectively, in *Arabidopsis* (Berthet *et al*., 2011). *Arabidopsis* mutant *lac4lac11lac17* lacks lignin in the root (Zhao *et al*., 2013). When complemented with *MsLAC1* (*Miscanthus sinensis*), the *Arabidopsis* double mutant *lac4 lac17* resulted in normal lignification. *Arabidopsis Atlac2* mutation study revealed its role in root lignification and root growth (Khandal *et al*., 2020), while *Atlac8* mutation showed a change in flowering time. *Atlac15* mutant plants showed change in the seed colour (Cai *et al*., 2006). *Atlac15* mutant plants exhibited an increase in the soluble proanthocyanidin or condensed tannin content in seeds (Liang *et al*., 2006a). Similar phenotype was found in the case of *BnTT10, PtLAC3*, and *ZmLAC3*, in B. napus, poplar, and maize, respectively (Zhang K *et al*., 2013; Ranocha *et al*., 2002; Caparros-Ruiz *et al*., 2006;). These enzymes are reported to be associated with abiotic stress regulation (Wei *et al*., 2000; Liang *et al*., 2006b; Turlapati *et al*., 2011; Shen *et al*., 2013). The overexpression of rice *LAC* gene *OsChI1* in *Arabidopsis* causes drought and salt stress tolerance in transgenic plants (Cho *et al*., 2014). *ZmLAC1 was induced* in maize primary roots when treated with varying concentrations of NaCl (Liang *et al*., 2006b). In *Cicer arietinum*, upregulation of *CaLaccase* genes along with increased lignin content was found in cold acclimated plants as compared to non-acclimated plants (Khaledian *et al*., 2015). *LACCASE* transcripts in maize were shown to be considerably elevated in response to Pb stress, which may play a role in Pb hyperaccumulation in roots (Shen *et al*., 2013). Cadmium stress increased the expression of FLA, a *LACCASE* involved in lignin synthesis and secondary cell wall deposition and thickening in cotton (Chen *et al*., 2019). *GhLAC1* and *GhLAC4* overexpression increases lignification in cotton, which leads to greater tolerance to the fungal pathogen *Verticillium dahliae* as well as the insect pests cotton bollworm (*Helicoverpa armigera*) and cotton aphid (*Aphis gosypii*) (Hu *et al*., 2018, Wei *et al*., 2021).

In *Arabidopsis*, under copper deficiency, miRNA negatively regulates several *LAC*s (Abdel-Ghany *et al*., 2008). Using prediction tools, At-miR408 has been predicted to target *LAC3, LAC12*, and *LAC13*. The *LAC2, LAC4*, and *LAC17* are predicted to be targeted by At-miR397 while, At-miR857 has been predicted to target the exon region of *LAC7* (Sunkar and Zhu, 2004; Abdel-Ghany *et al*., 2008). In *Arabidopsis*, lignin content and seed number are reported to be regulated by ath-miR397b (Wang *et al*., 2014). *Arabidopsis* miR397b was reported to regulate expression of *LAC2* and *LAC4* in the root and, thereby, promote lignin deposition in root under drought and low-phosphate (Khandal *et al*., 2020). The ptr-miR397a of poplar was predicted to target 29 *LAC* genes out of 49 (Lu *et al*., 2013). The overexpression of OsmiR397 in rice resulted in a higher yield. This miRNA expresses in young panicles and grains and negatively regulate *OsLAC* in rice. The *OsLAC13* (LOC_Os05g38420.1) regulated by OsmiR397 is involved in brassinosteroid sensitivity in rice (Jeong *et al*., 2011; Zhang Y *et al*., 2013). OsmiR397 imparts a domestication-related trait of low-lodging by increasing stem lignin deposition (Swetha *et al*., 2018).

In this study, we report that natural drought and dry root rot downregulate and upregulate the expressions of chickpea microRNA397 (CamiR397), and its targets *LAC4* and *LAC17L*, respectively. We demonstrate by overexpressing and inactivating CamiR397 that it regulates lignin deposition in root and, thereby, the responses towards drought and dry root rot in chickpea.

## Experimental Procedures

### Plant materials, growth conditions, construct preparation, Chickpea Transformation and Tissue Culture

Chickpea plants were grown in controlled conditions with a 22 °C temperature, 16 h photoperiod with a light intensity of 350 μmol m^−2^ s^−1^, and relative humidity of 70 %. The drought stress was imposed on the plants by withholding the water. For constitutive expression constructs of CamiR397 and STTM397, Gateway binary cloning technology (Invitrogen) was used according to manufacturer’s instructions. The precursor of CamiR397 and STTM construct was cloned in pGWB2 vector. The target mimic construct also known as **S**hort **T**andem **T**arget **M**imic is prepared according to Yan *et al*., 2012. A 48-base pair (core) adapter with antisense CamiR397 at its ends were prepared using PCR having three nucleotides (CTA) inserted at 10/11^th^ position in both antisense sequences. The CamiR397s in plants will bind to this transcript due to sequence complementarity but unable to cleave it since loss of cleavage specific site. Primer details are listed in Supplementary Table S3. For transformation, BGD72 variety of chickpea was used. The seeds were washed with MQ for 5 mins at normal shaking. Then the seeds were subjected to 70 % ethanol wash for 2 min at shaking. 2 % (v/v) Sodium hypochlorite was used to wash for 2-4 mins at shaking. After this, the seeds were washed thrice with MQ water at constant shaking. The seeds were now placed in water for 8 h for imbibe. The agrobacterium culture with desired construct was grown in LB broth containing 25 mg L^-1^ rifampicin and 50 mg L^-1^ kanamycin at constant shaking of 180 rpm in 28 °C incubator. Then this starter culture was used to grow secondary culture in LB broth with 50 mg L^-1^ kanamycin till the absorbance (at 600 nm) of 0.8. Then this culture was centrifuged at 2800 rpm for 5 mins. The obtained pellet was washed with half MS (Murashige and Skoog medium; Duchefa Biochemie, Netherlands) and resuspended in the half MS media. After resuspension, 0.5 ml of 100 mM acetosyringone was added and left for incubation at shaking for 30 mins.

Seed coats were removed from the imbibed seeds. After this, the seeds were cut in half along the embryo axis so that each cut part has half embryo. These wounded seeds were placed in culture flask containing resuspended culture and left for incubation at shaking for 30 mins. The seeds were removed and dried over sterilised filter paper. After removal of extra moisture, the seeds were transferred to half MS-plant agar plate for 48 h. The germinated seeds were transferred to full MS-agar plate with 50 mg L^-1^ kanamycin. After, every two weeks the regenerated shoots were transferred to a new MS-agar plate with increasing kanamycin concentration (100 mg L^-1^ and 150 mg L^-1^). The selected regenerated shoots were then micro-grafted to a wild type BGD72 plants. These grafted plants were grown under control condition (23 °C-25 °C and 60 % of humidity, 14 h dark period with a light intensity of 250 μmol m^−2^ s^−1^). *Agrobacterium tumefaciens* strain GV3101 was used for all transformations in chickpea.

### Soil moisture measurement and Relative water content

For receding water experiment, plants were grown in pot (60×25×25 LBH) in sand:soil mixture (1:1 v/v). These pots were well irrigated for 14 days after germination (dag). Irrigation was stopped in drought treated pots while continued irrigation in controlled pots. The drought stress level was measured using a soil moisture meter (Lutron Electronic Enterprise Co. Ltd., Taipei, Taiwan) at two depths (6 cm and 25 cm) at various spots in the pots for every alternate day. The soil moisture was maintained at 14-16 % in control pots while that of treated pots, decreased to as low as 4-7 % during the 14 days drought treatment.

### Lignin visualization and estimation

For the preparation of phloroglucinol stain solution, 2 gm of phloroglucinol powder (Sigma-Aldrich, USA) was dissolved in 80 ml of 20 % ethanol and 20 ml of concentrated HCl. The solution was kept in dark. The chickpea sections were immersed in the stain for 3-5 mins and washed once with distilled water very quickly. The sections were observed under Eclipse 80i (Nikon Instruments Inc., Japan) fluorescent microscope.

Acetyl bromide-based method was used to estimate the total lignin of root (Moreira-Vilar *et al*., 2014). The tissue was prepared to remove its protein and other UV-absorbing materials. This step was crucial in order to avoid measurement of other constituents at 280 nm apart from lignin. The tissue sample was harvested and freeze in liq. N_2_. After crushing in mortar-pestle with liq N_2_, the sample was homogenized with 7 ml of solution I (50 mM Potassium phosphate buffer-pH 7.0) and then transferred to a centrifuge tube. The homogenous mixture was centrifuged (1400xg, 5 min, RT). The pellet obtained was washed by sequential stirring and centrifugation onwards as follows: two times with sol I (50 mM Potassium phosphate buffer-pH 7.0), three times with sol II (1 % (v/v) Triton X-100 in sol I), two times with sol III (1 M NaCl in Sol I), two times with distilled water and two times with acetone. The washed pellet was dried in 60 °C oven for 24 h. After this, the pellet was vacuum cooled in desiccator. The dry pellet was defined as protein free cell wall extract. The protein free cell wall extract (20 mg) was incubated with 0.5 ml of 25 % acetyl bromide (v/v in glacial acetic acid) at dry bath of 70 °C for 30 mins. After incubation, the sample was quick chill in the ice. Then the sample was mixed with 0.9 ml of 2 M NaOH, 0.1 ml of 5 M hydroxylamine-HCl, and 5 ml of glacial acetic acid for the complete solubilization of lignin extract. Then the mixture was centrifuged at 1400xg for 5 min and the resultant supernatant was checked for absorbance at 280 nm. The absorptivity (□) was adopted from the report (Moreira *et al*., 2014) as 22.9 gm^-1^ L cm^-1^. The result was expressed as mg of lignin g^-1^ cell wall extract.

### GUS Reporter assay

1.5 kb promoter of miR397, *LAC4* and *LAC17L* were cloned upstream to GUS reporter gene in pGWB3 vector using Gateway system. For this, promoters were amplified from chickpea DNA through PCR using specific primers (Supplementary Table S3) and cloned in Gateway entry plasmid pENTR using D-TOPO cloning kit (Thermo Fisher Scientific) following manufacturer’s protocol. Subsequently, the insert was transferred to binary plasmid pGWB3 for the expression under CaMV35S promoter through LR clonase reaction (Thermo Fisher Scientific). Chickpea transformation was done with these constructs, as mentioned above. BGD72, after 2-3 weeks of growth, the plants were uprooted and checked for GUS activity. GUS activity was estimated by monitoring the cleavage of the substrate 4-methylumbelliferyl β-D-glucuronide (MUG) (Jefferson *et al*., 1987). The uprooted seedlings were washed with tap water and vacuum infiltrated with GUS staining solution (50 mM sodium phosphate buffer pH 7.0, 2 mM EDTA, 0.12 % Triton-X, 0.4 mM ferrocyanide, 0.4 mM ferricyanide, 1.0 mM 5-bromo-4-chloro-3-indoxyl-beta-D-glucuronide cyclohexyl ammonium salt (X-Gluc) for 20 mins and incubated in the dark at 37 °C overnight. The samples were cleared by incubating with a saturated solution of chloral hydrate at 65 °C for 1 h and visualized under light microscopy.

### Gas exchange parameters

At the vegetative stage, CO_2_ assimilation rate (A), stomatal conductance (g_s_), and transpiration rate (E) were measured using the leaf chamber fluorometer of an infrared gas analyzer (IRGA) (LI-6400/XT Portable Photosynthesis System, Lincoln, Nebraska, USA) with 350 µmol PAR, 400 µmol CO_2_, and 22 °C block temperature. Readings were logged five minutes after placing the chickpea leaf into the LCF chamber.

### MDA analysis

Lipid peroxidation levels were analysed by determining the MDA (malondialdehyde) production levels. The assay was done as described in (Pareek *et al*., 2019). About 200 mg of leaf tissue was taken, crushed in 5 % trichloroacetic acid (TCA), and centrifuged at 10,000□×□*g* for 10□min. Following this, an aliquot of the supernatant was added to a tube containing four parts of 0.5 % (v/v) thiobarbituric acid prepared in 20 % (v/v) TCA. These samples were heated at 95 °C for half an hour and then cooled down to 25 °C to terminate the reaction. The samples were centrifuged and absorbance for the supernatant was determined at 532□nm and 600□nm. The MDA content was then calculated using the following formula:

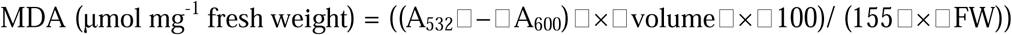

### Proline estimation

The proline content was estimated with colorimetric assay (Pareek *et al*., 2019). The 2^nd^ and 3^rd^ leaves (around 100 mg) from the top of chickpea plants were taken for estimation. 500 µL of 3 % Sulfosalicylic acid was added to homogenise liq N_2_ crushed samples. After centrifugation at RT for 5 min, 100 µL of supernatant was mixed with prepared reaction mixture (100□μL of 3 % sulfosalicylic acid, 200□μL glacial acetic acid, 200□μL acidic ninhydrin) and incubate at 96 °C for 1 h. The reaction was terminated after keeping it on ice. Then, 1 ml of toluene was added to this reaction and vortex extensively. After 5 min of incubation at RT, the phase with chromophore containing toluene was measured for absorbance at 520 nm.

### Sectioning with Cryostat microtome and vibratome

The tissues were harvested from 4-6 cm tap root zone from the root-shoot junction of the freshly uprooted roots. The cryo-microtome (CM1800) was used to cut transverse section after freezing the tissue in a freezing embedding medium at the tissue holder. The vibratome (Leica VT1000S) was used to cut the longitudinal sections of the root tissue. The samples were fixed with the help of 4-5 % agarose solution in a small tray. Then the embedded tissues were trimmed like a cube and placed on the tissue holder in vibratome. Sections were cut at the thickness of 50 µm.

### Infection of fungus

The blotting paper technique-based infection protocol described by Irulappan and Senthil-Kumar, 2021 was followed except for a fungal agar plug used for the infection. Agar plugs (∼7 mm diameter) with or without (Mock) *Macrophomina phaseolina* (ITCC 8635) from five days old culture plate were placed on roots and the plants were kept in a growth chamber with 25 °C ± 2 °C temperature, 16 h photoperiod with a light intensity of 150 μmol m^−2^ s^−1^, and relative humidity of 70 %. The observation was made 8 days after inoculation (DAI).

### *In-planta* fungal DNA quantification

The method described by Irulappan *et al*., 2021 was followed to estimate the fungal DNA present inside the root. Roots (∼1 cm long from infected areas) were collected at 8 DAI and quickly frozen in liquid N_2_ and stored at -80 °C. Total DNA was isolated and 50 ng was used to quantify using quantitative PCR. *M. phaseolina*-specific primers (Supplementary Table S3) were used to quantify the fungal load. A standard graph was prepared using a diluted known quantity of fungal DNA (100 ng, 10 ng, 1 ng, 0.1 ng, 0.01 ng, and 0.001 ng) which was used to calculate amount of fungal DNA in root samples.

### Microscopic observations, fluorescent staining, and Confocal microscope Imaging

Control and infected roots were observed for the presence of the necrotic lesion under a 0.5X objective lens of a stereomicroscope (SMZ25 Research Stereomicroscope; Nikon Corporation). Control and infected roots were collected and fixed in 10 % buffered formalin by vacuum infiltration. After 24 h, transverse sections (TS, 100 µm thickness) of roots were made using a vibratome (Leica VT1000 S; Leica Biosystems, Germany) and cleared with 10 % KOH overnight and neutralized with 1 M tris (pH 7.0). Then, TS sections were stained with wheat germ agglutinin-fluorescein isothiocyanate (WGA-FITC, Sigma-Aldrich) (maximum excitation, 495-500 nm, and maximum emission 514-521 nm), for fungal mycelia, and images were captured using Leica TCS SP5 Confocal (20X). TS sections were stained with basic fuchsin (Sigma-Aldrich) for staining lignin as described in the ClearSee protocol (Ursache *et al*., 2018). After staining, roots were captured under a 20X objective lens of a confocal microscope (SP5 or SP8) using 561 nm excitation and detection at 600–650 nm for basic fuchsin. The intensity of fluorescence µm^-2^ was measured using ImageJ (https://imagej.nih.gov/ij/) (Schneider *et al*., 2012).

### Scanning Electron Microscopy

For microscopic analysis, 50-micron sections of chickpea roots were fixed using 4 % (v/v) paraformaldehyde. The sample was placed on a metal stub, gold coated and then observed under a scanning electron microscope. The images were captured at a magnification of 1.0 K X and voltage of 20Kv. The images were analysed having a bar of 10 μm.

### Phylogenetic analysis

For phylogenetic analysis, full length protein sequence of *Medicago truncatula, Cicer arietinum* and *Arabidopsis thaliana* were aligned using the CLUSTALW algorithm. The aligned sequence was then used to generate a phylogenetic tree using the neighbor-joining method, using the “MEGA11” tool. The last two digits of predicted protein model accession numbers (XP) from NCBI are depicted for each species in the phylogeny. The list of protein accession numbers is presented in Supplementary Table S1. Bootstrap values are represented at each node, with a value of 100 indicating 100% support.

### RNA isolation and Expression analysis

Total RNA was extracted using TRI reagent (Sigma-Aldrich, USA) according to manufacturer’s protocol. For miRNAs, stem-loop primers were designed and cDNA for individual miRNA was prepared by pulsed-RT reaction using SuperScript III Reverse Transcriptase (Thermo Fisher Scientific) (Chen *et al*., 2005; Varkonyi-Gasic *et al*., 2007). Target gene expression analysis was done by quantitative RT-PCR (qRT-PCR). Primers used in this study are provided in Supplementary Table S3. For chickpea laccases, the primers for qRT-PCR are listed with the name followed by last two digits of XM accession number provided in Supplementary Table S1. qRT-PCR experiments were performed in Vii A7 Real-Time PCR System (Applied Biosystems CA, USA) using three technical and three biological replicates. Relative expressions of genes were calculated according to delta-deltaCt method of the system (Livak and Schmittgen, 2001). *CaEF-1*α (NM_001365163.1) (*elongation factor 1-*α) gene was used as internal control to normalize the variation in amount of cDNA template in chickpea samples (Garg *et al*., 2010).

## Results

### Drought increases Lignin deposition in chickpea root

Most studies apply drought to plants using mannitol or polyethylene glycol. This artificial treatment quickly causes uniform dehydration of the tissues dipped in the solution. In contrast, natural drought in the soil is a slow and gradual process, and the moisture content varies in gradient with the depth of the soil. To understand the drought-induced changes in chickpea physiology and phenotype, drought was applied to chickpea seedlings by withholding irrigation to the soil to mimic the natural drought condition. Pot-grown drought tolerant chickpea variety BGD72 (Sachdeva *et al*., 2020) was used for this experiment. After germination, the plants were grown with normal irrigation for 14 days. Soil moisture in the pot was measured at two different depths (6 cm and 25 cm) at various spots in the pots. The upper soil layers had 14-15 % relative soil moisture content, while the lower layers had 16-17 % during normal irrigation of pots. Irrigation was stopped from the 15^th^ day onwards to impose drought, while irrigation to the control plants continued. After 28 days after germination (dag), the relative soil moisture content decreased to ∼4 % and ∼7.5 % at the upper and lower layers, respectively, in the drought-treated pots (Supplementary Fig. S1A). Root length was measured for both the control and drought-treated plants after the completion of treatment on 28^th^ dag. The drought-treated plants exhibited ∼28 % longer tap roots compared to the well irrigated control plants. The roots appear thinner with fewer lateral roots (Fig. 1A-B.). Drought- or salinity-induced lignin deposition in the root vascular tissues has been reported in crop plants (Fan *et al*., 2006; Jbir *et al*., 2001). Phloroglucinol staining of the longitudinal sections of the root zone 5-6 cm beneath the root-shoot junction displayed increased lignin deposition in drought-treated samples (Fig. 1C). Estimation of lignin content of the whole primary root by acetyl bromide method (Moreira-Vilar *et al*., 2014) showed about 27 % increase in lignin in chickpea var. BGD72 roots (Fig. 1D). The transverse section of the same region showed lignin deposition predominantly in the xylem vessels, ray, and parenchyma within the vascular bundle (Fig. 1C). Lignin estimation in the plant’s stem tissue exhibited no significant change in lignin content in this cultivar under the same condition (Supplementary Fig. S1B).

**Figure 1.**
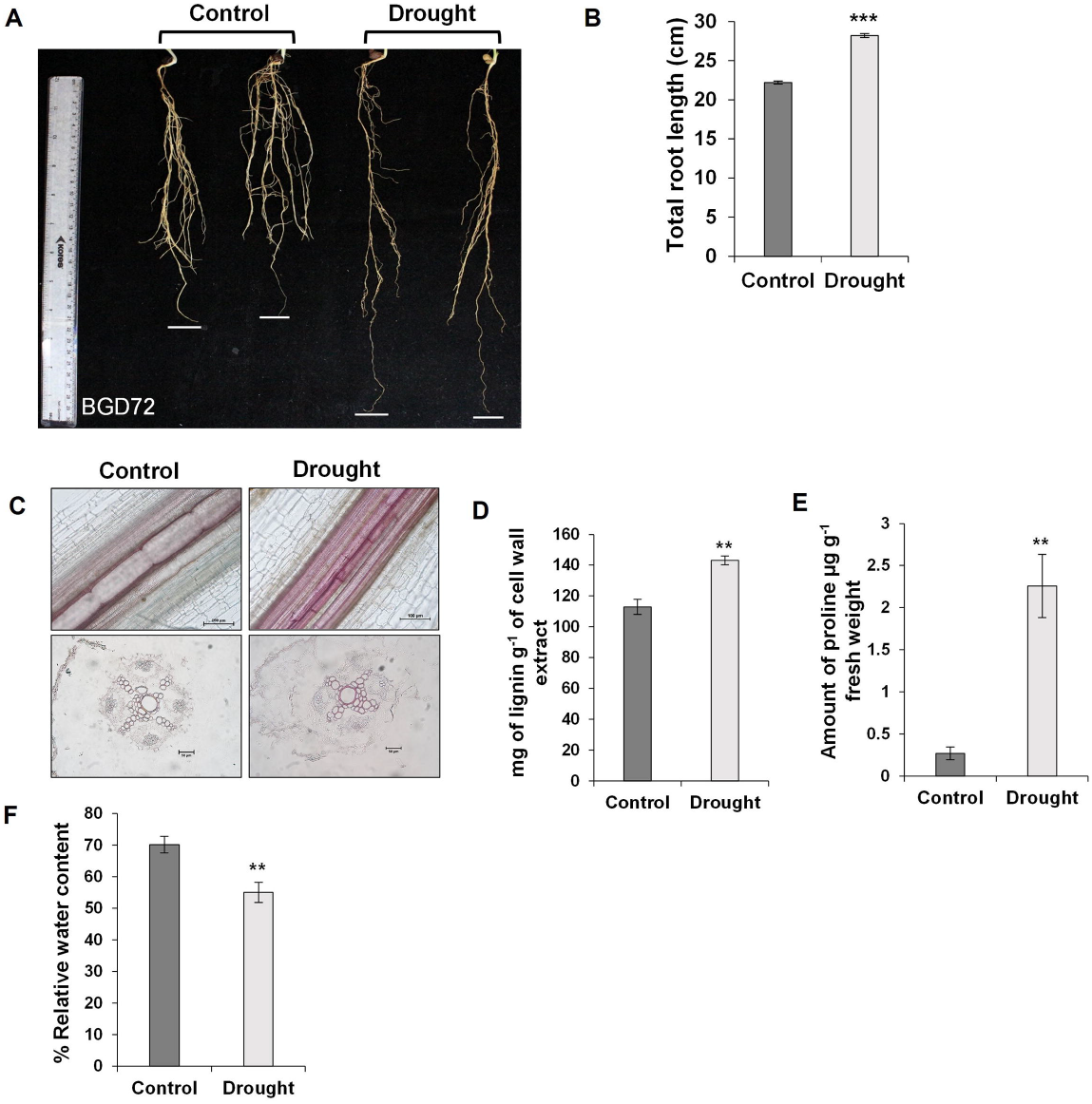
Morphological, physiological changes and lignin deposition in chickpea roots under drought stress. **A**. Pictorial illustration of chickpea cultivars (BGD72) roots under drought stress. Chickpea plants were grown under control and drought conditions as described Sharma *et al*., 2020. **B**. Measurement of primary root length in chickpea under control and drought-exposed condition (n = 31). The measurements were taken at the end of the experiment. **C**. Transverse sections (TS) of primary roots under control and drought conditions in the chickpea cultivars. Sections were taken from the root at 6–7 cm below the root-shoot junction. Lignin deposition was shown by phloroglucinol staining in xylem tissue of the root. **D**. Total lignin content was estimated in control and drought-exposed chickpea primary roots. Lignin content was estimated by acetyl bromide method and presented as mg of lignin g^-1^ of cell wall extract. **E**. Proline content (in µg g^-1^ fresh weight) under control and drought conditions. Proline estimation was done with Colorimetric assay. **F**. Relative water content (RWC) in third and fourth leaves from top of chickpea under control and under drought-treated conditions. Plants were grown 14 days with normal hydration. Samples were harvested for analysis after 14 days of receding water condition. Asterisks indicate significant differences from the control as determined by Student’s *t*-test (* for *p* < .05; ** for *p* < .01; *** for *p* < .001). Error bars show ± SE.

Imposition of the drought was positively correlated with 8-fold higher proline content in the third and second leaves from the top, and a 15 % decrease in the leaf relative water content (RWC) in the third and fourth leaves from the top of the plants (Fig. 1E-F), and the increased expression of drought inducible gene, 9-*cis-EPOXICAROTENOID DIOXYGENASE* (*CaNCED2*; XM_004504855) in the drought-treated plants (Supplementary Fig. S1C).

### *LACCASE* and MicroRNA expression profile in the chickpea root under drought

A search in the published annotated chickpea genomes and transcriptome (Varshney *et al*., 2013; Jain *et al*., 2013, Garg *et al*., 2011) led to the identification of 20 LACCASE-encoding genes and transcripts. Expression profiles of the *LAC* genes were monitored in the control and drought-treated roots, and 16 *LACCASEs* showed expression in the chickpea root under drought through qRT-PCR (Fig. 2A). A small RNA sequencing of the control and drought-treated chickpea root resulted in 15 differentially expressed microRNAs in the drought-treated samples with nine upregulated and six downregulated microRNAs. A heatmap showing the expression profiles of the increased and decreased well-known miRNAs in the drought-treated chickpea roots (Fig. 2B). Out of those, interestingly, a miRNA (CamiR397) showing homology to miR397 of *Arabidopsis* was found downregulated under drought. MiR397 was predicted to target *LAC* transcripts, which regulate lignin deposition in plants (Sunkar and Zhu, 2004; Abdel-Ghany *et al*., 2008; Lu *et al*., 2013). MicroRNA397 was also downregulated in rice in response to drought (Bakhshi *et al*., 2016). However, it contrasts the upregulation of miR397b in response to mannitol treatment in *Arabidopsis* (Khandal *et al*., 2020). Unlike in *Arabidopsis*, we found only one CamiR397 species in the previously sequenced chickpea small RNA libraries (Jain *et al*., 2014), and CamiR397 showed sequence similarity with miR397s of other plant species (*Medicago truncatula, Glycin max, Arabidopsis thaliana*) (Fig. 2C). The precursor sequence of CamiR397 was derived from the chickpea genome sequence (Varshney *et al*., 2013). Based on the reported standards by Zuker, 2003, the secondary structure predicted for the precursor of CamiR397 represents the location of mature micro-RNA sequence on hairpin structure of the precursor stem-loop (Fig. 2D). The mature sequence of miR397 in the predicted secondary structure were present with mismatches for hairpin recognition by pre-miRNA processing enzymes with free energy (ΔG) of -43.90 kJ mol^-1^.

**Figure 2.**
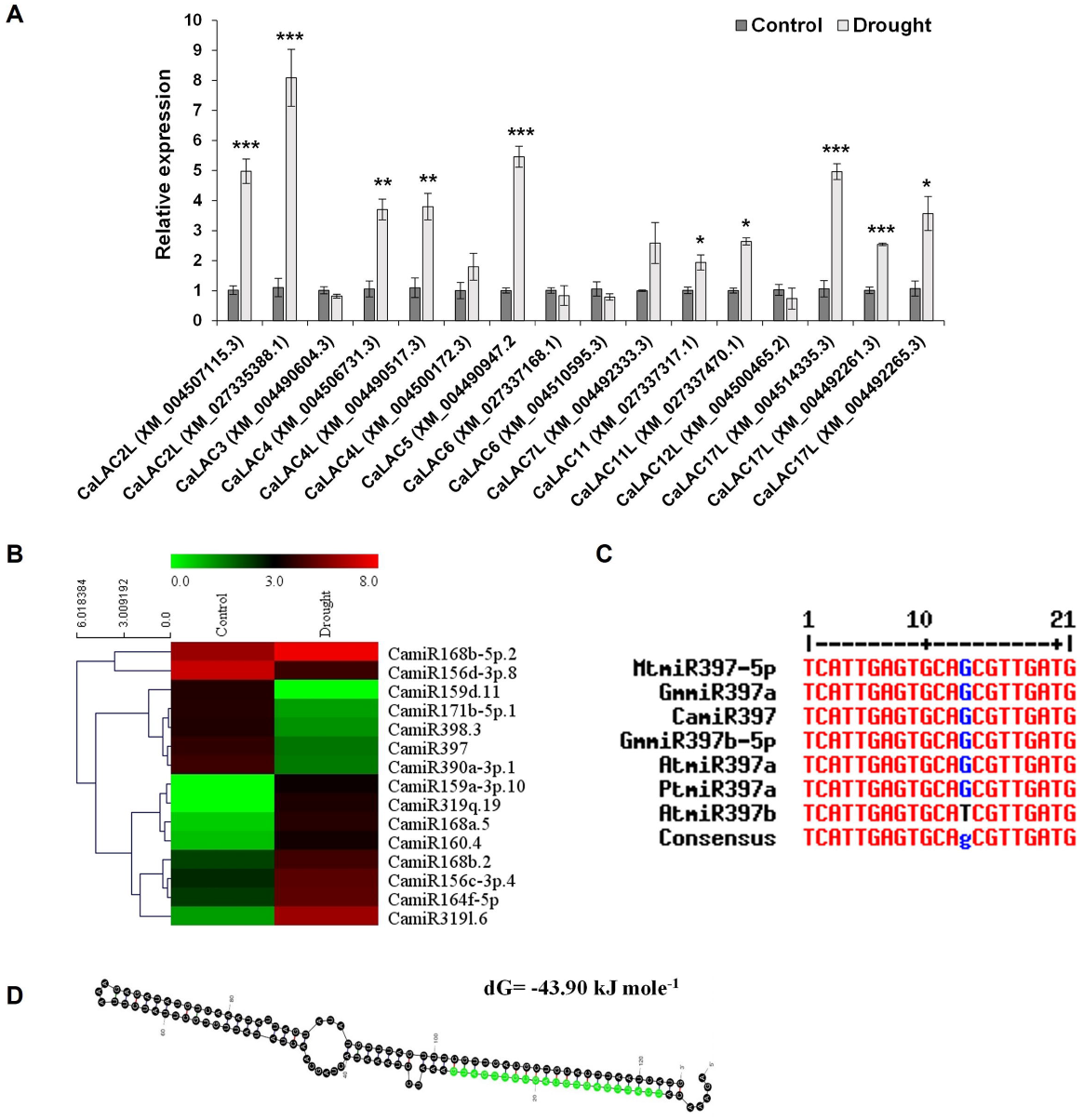
LACCASE expression profile in Chickpea and CamiR397 expression heat map. **A**. Relative expression profile of chickpea *LACCASEs* under control and drought condition. The laccases are labeled with their predicted mRNA model ID (XM in NCBI) (Supplementary Table S1). BGD72 plants were grown 14 days with standard irrigation. Irrigation was stopped in drought at 15^th^ day till the end of experiment i.e., 28^th^ day, while irrigation was continuous in control. Samples were harvested for analysis after 14 days of receding water condition. *CaEF1*α (NM_001365163.1) was taken as internal control (Garg *et al*., 2010). **B**. Heat map for differentially expressed miRNAs identified in RNAseq data generated from libraries of chickpea cultivar BGD72 under control and drought treatment post small RNA sequencing. Heat map is based on their normalized reads per million (RPM) values. **C**. Sequence alignment of mature miRNA397 of different plant species viz. *Medicago truncatula* (Mt), *Glycine max* (Gm), *Cicer arietinum* (Ca), *Arabidopsis thaliana* (At) and *Populus trichocarpa* (Pt) using Multialin web portal. **D**. Predicted precursor hairpin structure of CamiR397 formed after cleavage of primary miRNA. Asterisks indicate significant differences from the control as determined by Student’s *t*-test (* for *p* < .05; ** for *p* < .01; *** for *p* < .001). Error bar show ± SE.

### Expression of CamiR397 and its predicted targets in chickpea

A phylogenetic analysis of the full-length LACCASE proteins of *Medicago truncatula, Arabidopsis thaliana*, and chickpea (*Cicer arietinum*) distributed the proteins into six clades (Fig. 3). All chickpea LACCASEs (Taxon names of all LAC proteins are associated with last two decimals of XP id in NCBI, and listed in Supplementary Table S1) were dispersed throughout the phylogenetic tree, which suggests relatively low sequence homology. Chickpea LACCASE proteins were named according to sequence similarity with their *Arabidopsis* orthologs. Using a small RNA target prediction algorithm, the predicted targets for CamiR397 were the mRNAs of *LACCASE 4* (*LAC4*) (XM_004506731.3), *LACCASE 4 LIKE* (*LAC4L*) (XM_004500172.3) and *LACCASE 17 LIKE* (*Lac17L*) (XM_004514335.3) (Supplementary Table S2). In the phylogeny tree, groups II and III consist of the target *LACCASEs* of CamiR397 (Fig. 3). The predicted target sequences showed alignment with CamiR397 with perfect and imperfect match (Fig. 4A). Expression of CamiR397 and the predicted mRNA targets were assessed through stem-loop qRT-PCR (SLqRT-PCR) and qRT-PCR under control and drought. The expression of CamiR397 was decreased in the drought-treated sample by almost 3-fold, and the targets, *LAC4* and *LAC17L*, showed 4.3- and 3.3-fold increase in the expression under drought, while *LAC4L* didn’t show any significant change under drought (Fig. 4B-C). This is in contrast to miR397b expression in *Arabidopsis*, in which miR397b was found downregulated, and miR397a expression level was unchanged under mannitol treatment for 24 hours (Khandal *et al*., 2020; Sunkar and Zhu, 2004). The inverse correlation indicated the target-miRNA relationship between *LAC4* and *LAC17L*, and CamiR397.

**Figure 3.**
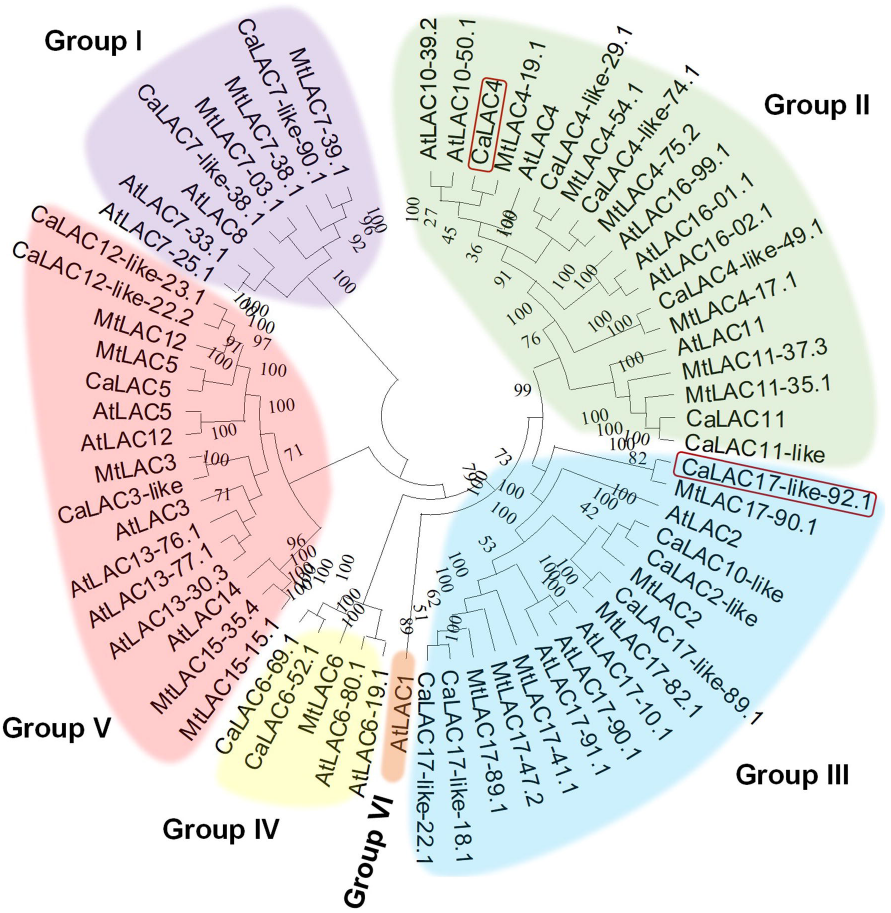
Phylogenetic tree of all laccases of Chickpea, Medicago and Arabidopsis. Neighbor-joining tree of protein sequences of 20 LACCASES of *C. arietinum*, 21 LACCASES of *M. truncatula* and 17 LACCASES of *A. thaliana*. The last two digits of predicted protein model accession numbers (XP in NCBI) are depicted for each species in the phylogeny. The list of protein accession numbers are presented in Supplementary Table S1. The abbreviations are representative laccases of three species. CaLAC-*Cicer arietinum* LACCASE, MtLAC-*Medicago truncatula* LACCASE and AtLAC-*Arabidopsis thaliana* LACCASE. Bootstrap values are represented at each node, with a value of 100 indicating 100% support.

**Figure 4.**
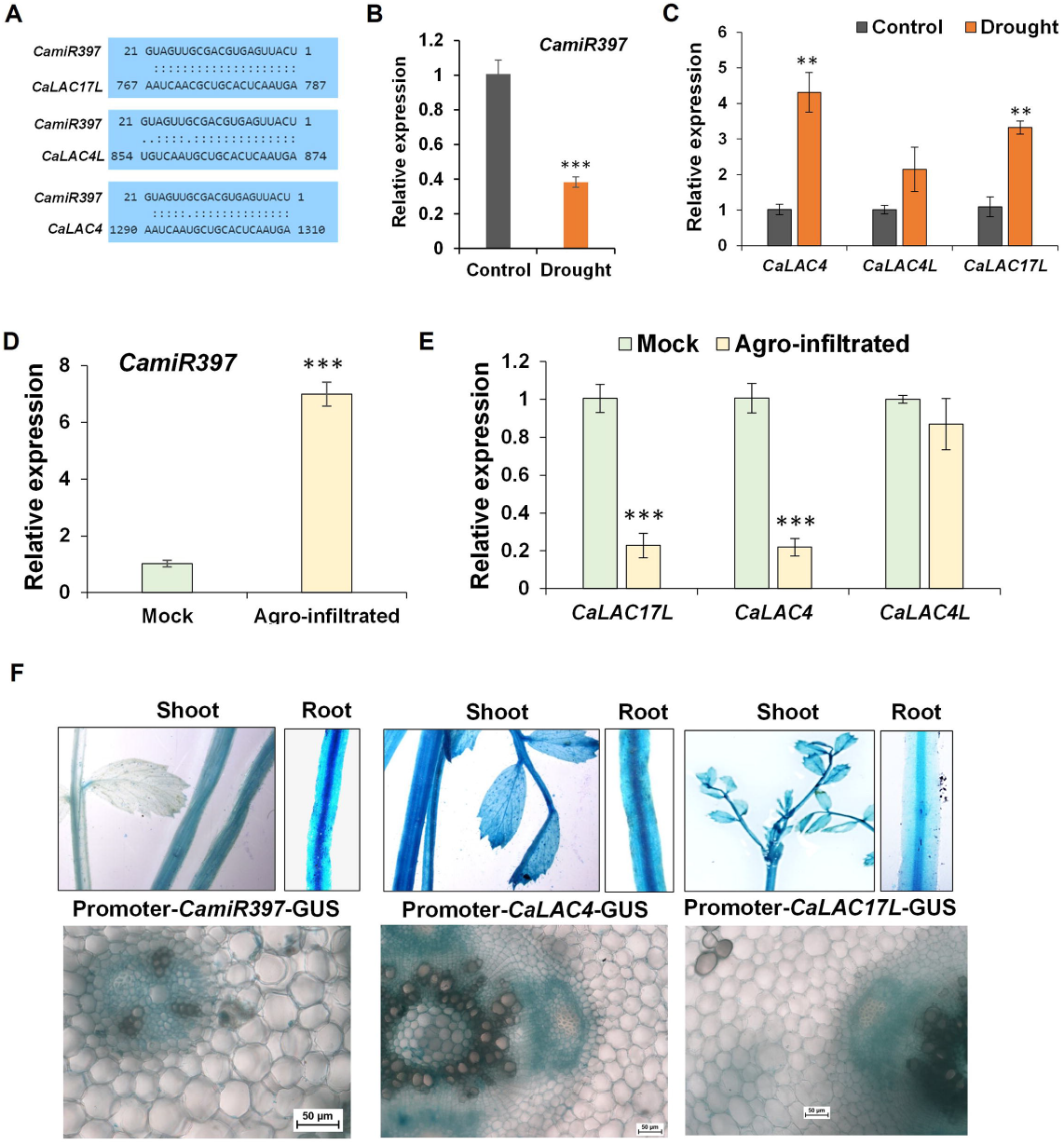
Expression profile of CamiR397 and predicted targets. **A**. Predicted target sequences of CamiR397. **B** & **C** Expression pattern of the CamiR397 and their predicted target genes in response to the drought condition as assessed by qRT-PCR. Asterisks indicate significant differences from the control as determined by Student’s *t*-test (* for *p* < .05; ** for *p* < .01; *** for *p* < .001). **D** & **E** CamiR397-target interaction was validated by transient over-expression of miRNA precursor sequence in chickpea leaf through agroinfiltration and change in expression pattern of their respective target genes (*CaLAC4, CaLAC4-like* and *CaLAC17-like*) were assessed by qRT-PCR as compared with control sample. Three replicates were taken. *CaEF1*α (NM_001365163.1) was used as internal control for normalization. Asterisks indicate significant differences from the mock as determined by Student’s *t*-test (* for *p* < .05; ** for *p* < .01; *** for *p* < .001). **F**. Tissue-specific expression and localization of GUS activity in various tissues of CamiR397 and their targets (*pCamiR397::GUS, pCaLAC4::GUS* and *pCaLAC17L::GUS*) expressing transgenic Chickpea lines. Transverse section of roots of GUS expressing transgenic lines showing GUS activity in various cell types. Error bars show ± SE.

To establish the target-miRNA relationship between *LAC4* and *LAC17L* genes with CamiR397, the precursor of CamiR397 was cloned under CaMV35S promoter and was expressed transiently in chickpea leaves by agroinfiltration. CamiR397 expression was increased by 7-fold in all the biological replicates. In contrast, the expression of its target genes LAC4 and LAC17L showed a 5-fold reduction compared to the sample infiltrated with *Agrobacterium* containing empty vector (mock). In contrast, the reduction in *LAC4L* expression was not significant, suggesting *LAC4* and *LAC17L* mRNAs are targets of CamiR397 (Fig. 4D-E). RNA ligase-mediated rapid amplification of cDNA ends (RLM-RACE) followed by sequencing validated cleavage of *LAC4* and *LAC17L* mRNAs at the predicted *miR397b* target sites (673^rd^ and 694^th^ nucleotides, respectively, from the translation initiation site) (Supplementary Figure Fig. S1D).

To investigate the expression of CamiR397 and its target genes in the same tissue, the 1.5 kb-long upstream sequences (Supplementary Text S1) of CamiR397, *LAC4*, and *LAC17L* genes were fused with GUS (β-Glucuronidase) gene and introduced into chickpea var. BGD72 for stable transformation. Transgenic T2 seedlings were stained for GUS activity. A strong GUS activity was observed in the whole seedlings, including the root in the *pCaLAC4:GUS-* and *pCaLAC17L: GUS-*expressing plants, although the GUS activity was higher in the case of the *LAC4* promoter. The *pCamiR397:GUS* plants did not show GUS expression in the leaves and showed a very weak expression in the stem. In these plants, the GUS activity was primarily observed in the root vascular tissue and cortex (Fig. 4F), showing that CamiR397 and its target genes are expressed in the same tissue.

### CamiR397 regulates lignin deposition in chickpea roots and response towards drought

To investigate the role of CamiR397 in root lignin deposition, stable transgenic chickpea plants overexpressing CamiR397 (miR397OX) were developed using a CaMV35S promoter fused to its precursor sequence. To analyse the effect of biological inactivation of CamiR397, a short tandem target mimic molecule (STTM397) was synthesized following Yan *et al*., 2012 (Supplementary Fig. S2), and the overexpressing lines of the STTM397 constructs were developed. Two independent representative T3 transgenic lines for each construct were used for further studies. The relative expression of CamiR397 in miR397OX lines exhibited an increased expression of the miRNA by 2.5-3.5-fold under both control and drought conditions. In contrast, STTM397 lines expectedly showed decreased expression of CamiR397 under drought (Fig. 5A). Both the CamiR397-overexpressing transgenic lines showed 2 to 2.4-fold reduced expression of target genes *LAC4* and *LAC17L*, in the control and drought conditions. The target mimic lines exhibited 2.5 to 3-fold higher expression of the target genes under both conditions (Fig. 5B-C). Expression levels of other *LAC* genes in the CamiR397 and STTM397 lines showed no significant change compared to the wild type (Supplementary Fig. S3).

**Figure 5.**
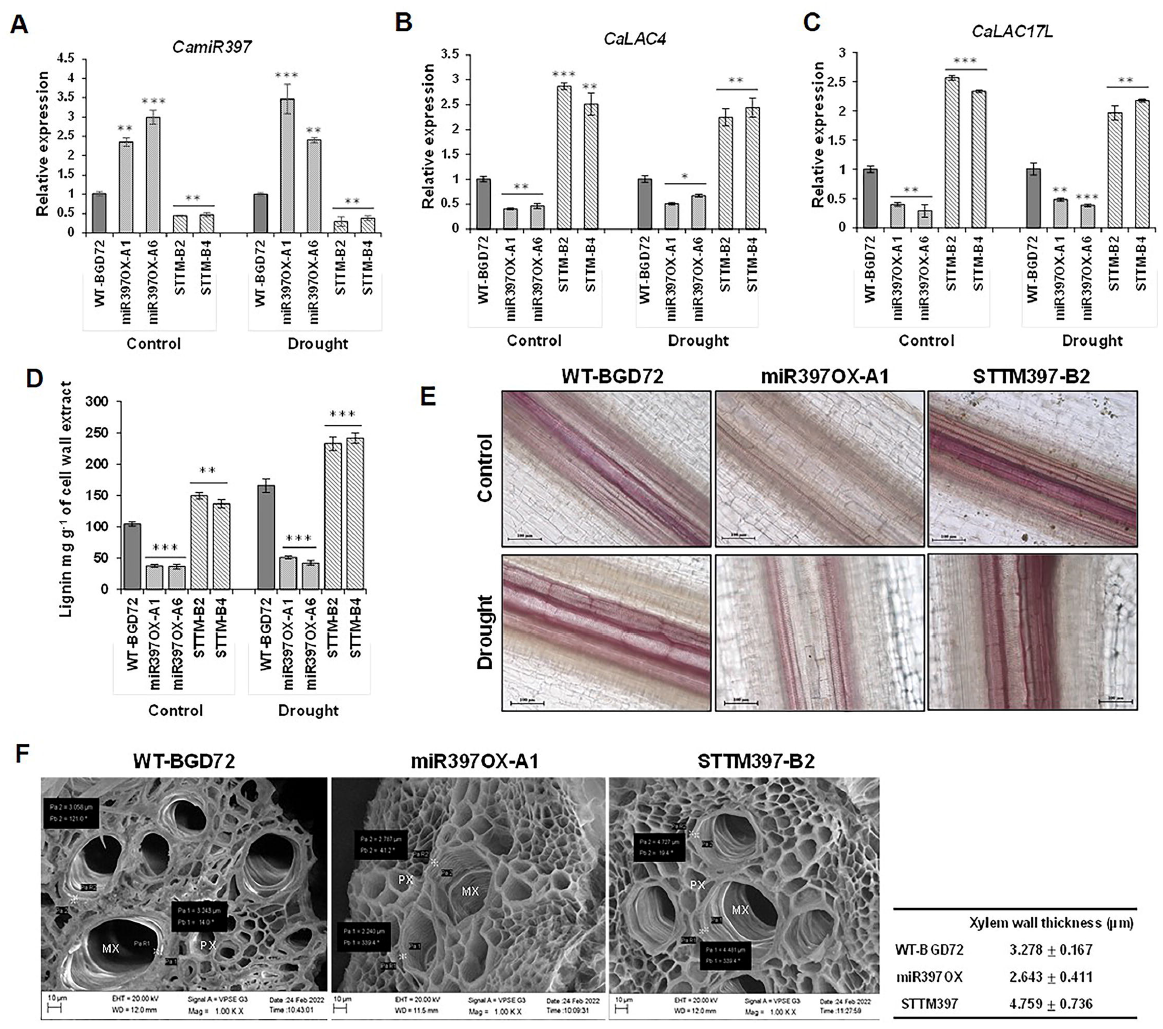
CamiR397-Target expression profile and Lignin deposition in the root of wild type and transgenic chickpea lines under control and drought stress condition. **A**. Relative expression of CamiR397 in miR397OX (A1 & A6) and STTM397 (B2 & B4) lines compared with wild type BGD72 plants (WT-BGD72) under control and drought condition as assessed by stem loop qRT-PCR. **B** & **C** Relative expression of *CaLAC4* and *CaLAC17L* genes in miR397OX and STTM397 lines compared with WT under control and drought condition as assessed by qRT-PCR. **D**. Lignin content estimation in miR397OX and STTM397 lines compared with WT plants under control and drought condition through acetyl bromide method. The unit of lignin measurement was presented as mg of lignin g^-1^ of cell wall extract. Two transgenic lines with three replicates were selected for the study. **E**. Phloroglucinol staining in the longitudinal sections of root xylem tissue of 28 days old WT, miR397OX and STTM397 lines under control and drought condition. Sections were taken from the root at 6–7 cm below the root-shoot junction. Images were taken at 20X magnification to show the lignin deposition. Scale bars, 100 μm. **F**. Scanning electron microscope micrographs of transverse sections through root xylem of WT, miR397OX (A1) and STTM397 (B2). The transverse section of root exhibiting protoxylem (PX) and metaxylem (MX) vessels with secondary wall thickening and pits distributed in non-uniform manner. Filled boxes showed thickness of xylem wall presented with micrometer (µm). Table to represent change in xylem wall thickness in wild type and transgenic plants using measurement tools in SEM. Asterisks indicate significant differences from the WT-BGD72 as determined by Student’s *t*-test (* for *p* < .05; ** for *p* < .01; *** for *p* < .001). Error bars show ± SE.

Root lignin content was measured in these lines. The miR397OX lines showed 2.5-fold lower lignin content, while the STTM397 lines exhibited 1.5-fold higher lignin content (Fig. 5D).To investigate the effect of an increase in the root lignin content in the drought response, plants were subjected to drought treatment as mentioned above. The control line showed a 1.5-fold increase in the root lignin. There was a marginal increase in the root lignin content in the CamiR397-overexpressing line under drought over the same line, while the root lignin content in the STTM397 lines increased ∼1.3-1.5-folds over the same lines under the control condition (Fig. 5D). This observation suggested that LAC proteins other than LAC4 and LAC17L might also have roles in the lignin deposition. These results were supported by phloroglucinol staining of the root longitudinal sections (Fig. 5E). Transverse sections of the roots of both the wild type (WT-BGD72) and transgenic lines (miR397OX and STTM397) were analyzed under the scanning electron microscope. The miR397OX lines showed a thinner (∼20 % decrease) xylem wall, and the STTM397 lines exhibited a ∼45 % thicker xylem wall compared to the non-transformed plants (Fig. 5F).

To examine the biological significance of increased and decreased root lignin deposition, we assessed the growth and photosynthetic parameters of the transgenic lines. The transgenic plants did not show any significant deviation in the vegetative and reproductive parameters from the non-transformed plants as measured by plant height, internodal distance, and seed weight (Supplementary Fig. S4). However, the photosynthetic ability differed from the non-transformed plants as measured by CO_2_ assimilation rate, transpiration rate, and stomatal conductance from the third and fourth leaf from the top of the plants exhibited change under drought stress in WT and transgenic lines. There was no significant change in these parameters between the WT and transgenic lines under control conditions. The overexpression lines showed decreased assimilation rate and transpiration rate as compared to WT under drought condition, whereas STTM397 lines exhibited significantly improved CO_2_ assimilation rate, transpiration rate, and stomatal conductance under drought compared to WT plants (Fig. 6). The overexpression lines exhibited ∼35-50 % decrease in assimilation rate under drought compared to WT, while only one overexpression line showed a significant reduction of ∼50 % in transpiration rate. The STTM397 lines showed a slightly improved assimilation rate showing an increase of about 30 % from the wild type, while stomatal conductance under drought conditions increased ∼25 %. While one STTM line showed an improved transpiration rate of about 20 % under drought, the increase in transpiration rate in the other line was not significant (Fig. 6A, B-C).

**Figure 6.**
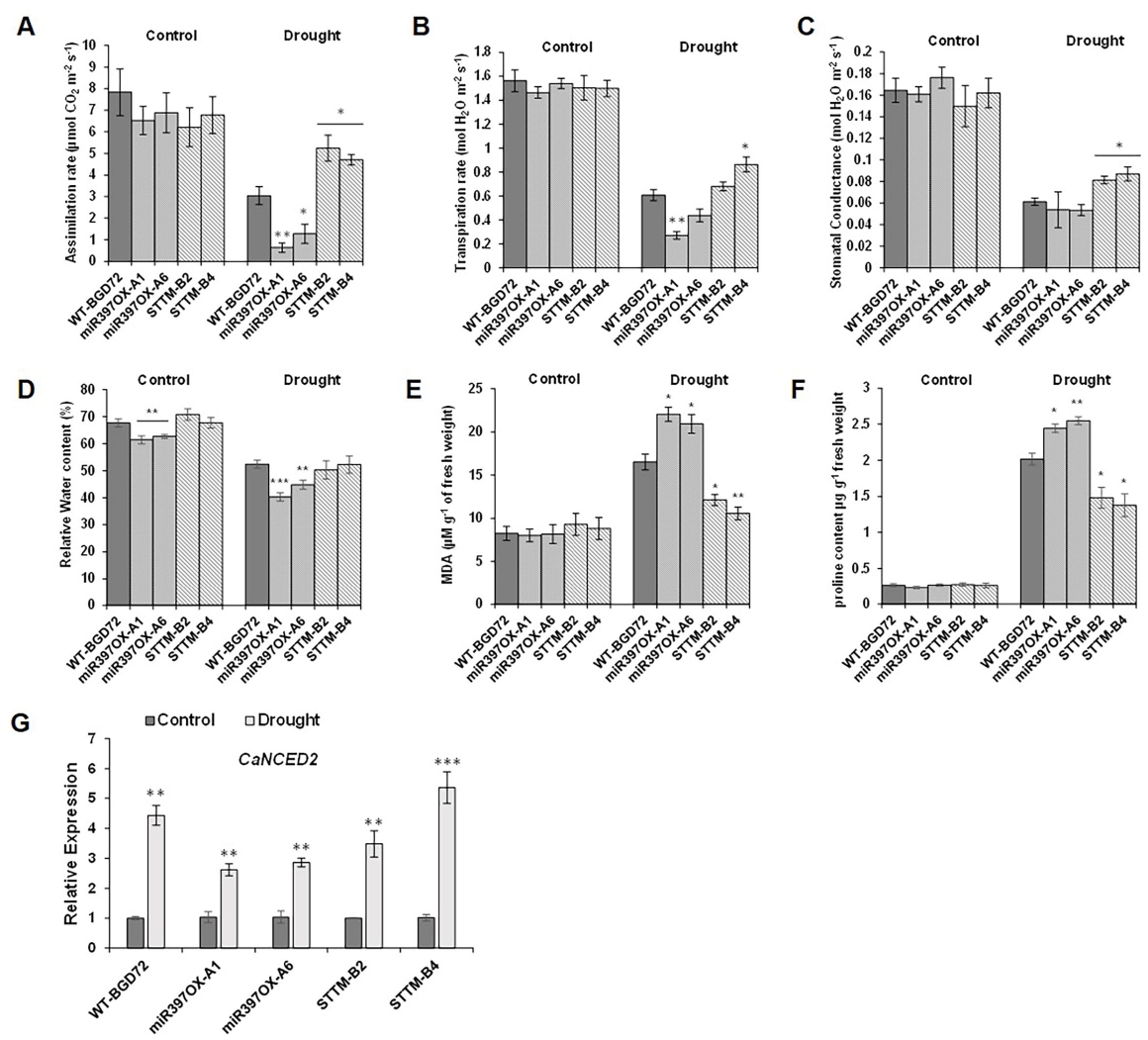
Evaluation of environmental stress physiology in transgenic lines and wild type plants. **A, B** & **C** Measurement of agronomic parameters (Assimilation rate, Stomatal conductance and Transpiration rate, respectively) in miR397OX (A1 & A6) and STTM397 (B2 & B4) lines compared with WT-BGD72 under control and drought condition (n=5). Plants were grown and given drought treatment as mentioned above. **D**. Relative water content (RWC) in third and fourth leaves from top of both transgenic and WT plants at control and drought stress conditions. **E** & **F** Malondialdehyde (MDA) content and Proline content were measured in WT and transgenic lines under control and drought conditions. Proline estimation (µg g^-1^ fresh weight) was done as previously mentioned. MDA content (µM g^-1^ fresh weight) was estimated through thiobarbituric acid (TBA) assay. **G**. Relative expression level of ABA synthesis gene, 9-*cis-EPOXYCAROTENOID DIOXYGENASE* (*CaNCED2*; XM_004504855) in the root of WT and transgenic lines under control and drought condition. Asterisks indicate significant differences from the WT-BGD72 as determined by Student’s *t*-test (* for *p* < .05; ** for *p* < .01; *** for *p* < .001). Error bars show ± SE.

Drought-response parameters such as RWC, proline, and MDA (Malondialdehyde). RWC was measured in WT and transgenic plants after 14 days of drought treatment. The miR397OX lines showed an 8-10 % reduction in leaf water content under normal conditions. All the plants showed a 25-30 % decrease in RWC under drought. The miR397OX lines showed a 15-20% reduction in RWC as compared to the control plants under drought, while the STTM397 lines didn’t show any significant difference (Fig. 6D). Similar MDA contents were observed in all the plants in the control condition. However, under drought, an increase and decrease of ∼25% in MDA content as compared to the control plants were observed for the miR397OX and STTM397, respectively (Fig. 6E). Under drought conditions, an 18-20 % increase in proline accumulation was found in the miR397OX lines, while the STTM397 lines exhibited 20-25 % reduction in proline content in comparison to WT (Fig. 6F). The expression of ABA synthesis gene, 9-*cis-EPOXYCAROTENOID DIOXYGENASE*(*CaNCED2*; XM_004504855) was increased in all plants under drought condition (Fig. 6G).

### CamiR397 can regulate tolerance to dry root rot disease in chickpea

Plants form barriers in specific cell types to combat adverse environment. Endodermal Casparian strips made of lignin and suberin constitute a well-known barrier in the roots. Exodermis or hypodermis, located just under the outermost cell layer epidermis, also forms a lignified barrier in some angiosperms. This is thought to allow the plant to retain water in drought (Peterson and Perumalla, 1990). We checked for any lignin deposition in the exodermis of chickpea root by staining with a fluorescent dye basic fuchsin. Transverse sections of soilrite-grown chickpea roots showed less but clear lignin deposition in the exodermis, endodermis, and vascular tissues (Supplementary Fig. S5A). The lignin deposition in the root exodermis layer in the wild type, miR397OX, and STTM397 lines followed their relative values of root lignin content assay (Supplementary Fig. S5B).

Dry root rot (DRR) caused by the fungus *M. phaseolina* is an economically important disease in chickpea and can cause 3-70 % yield loss in India and other sub-tropical chickpea-growing countries (Nene and Sheila, 1996; Assfaw and Negash, 2020; Rai *et al*., 2022). So far, molecular identification of any resistant gene has not been reported, and application of fungicides in the seeds before sowing is followed as the only management of the disease control. Infection with *M. phaseolina* induced expression of *LAC3* and *MYB* involved in the lignin biosynthesis pathway (Irulappan *et al*., 2022). We inoculated *M. phaseolina* in chickpea root to monitor the plant’s response in terms of *LACCASE* gene expression. The fungal infection decreased CamiR397 expression in root by ∼2-fold. Simultaneously, expression of nine chickpea LAC genes were increased (Fig. 7A). The lignin deposition reflected the increase in LAC gene expression in the exodermis and around the infection site (Supplementary Fig. S6). In order to investigate any effect of the variable lignin deposition in the exodermis of the transgenic chickpea lines in the infectivity of *M. phaseolina*, we compared the lesion length, the number of cortical layers colonized by fungus, and the fungal DNA accumulation as the extent of infection. The WT plant showed a moderate infection and accumulation of lignin in the exodermis and around the site of infection (Fig. 7B, C-D). The miR397OX chickpea roots were highly sensitive to infection and exhibited much-reduced lignin accumulation. In contrast, STTM397 plants showed high resistance against the infection and enhanced lignin deposition in the exodermis and around the site of the infection. The relative infection level was also monitored by fluorescence staining of fungal hyphae and fungal DNA quantification. The miR397OX lines had more fungal colonization than the STTM397 lines (Fig. 7E, F-G). Our results suggest that local lignin deposition in the root exodermis and around the infected region determines the extent of resistance against dry root rot disease in chickpea and CamiR397 can regulate the lignin deposition in the exodermis and, thereby, the resistance against *M. phaseolina*.

**Figure 7.**
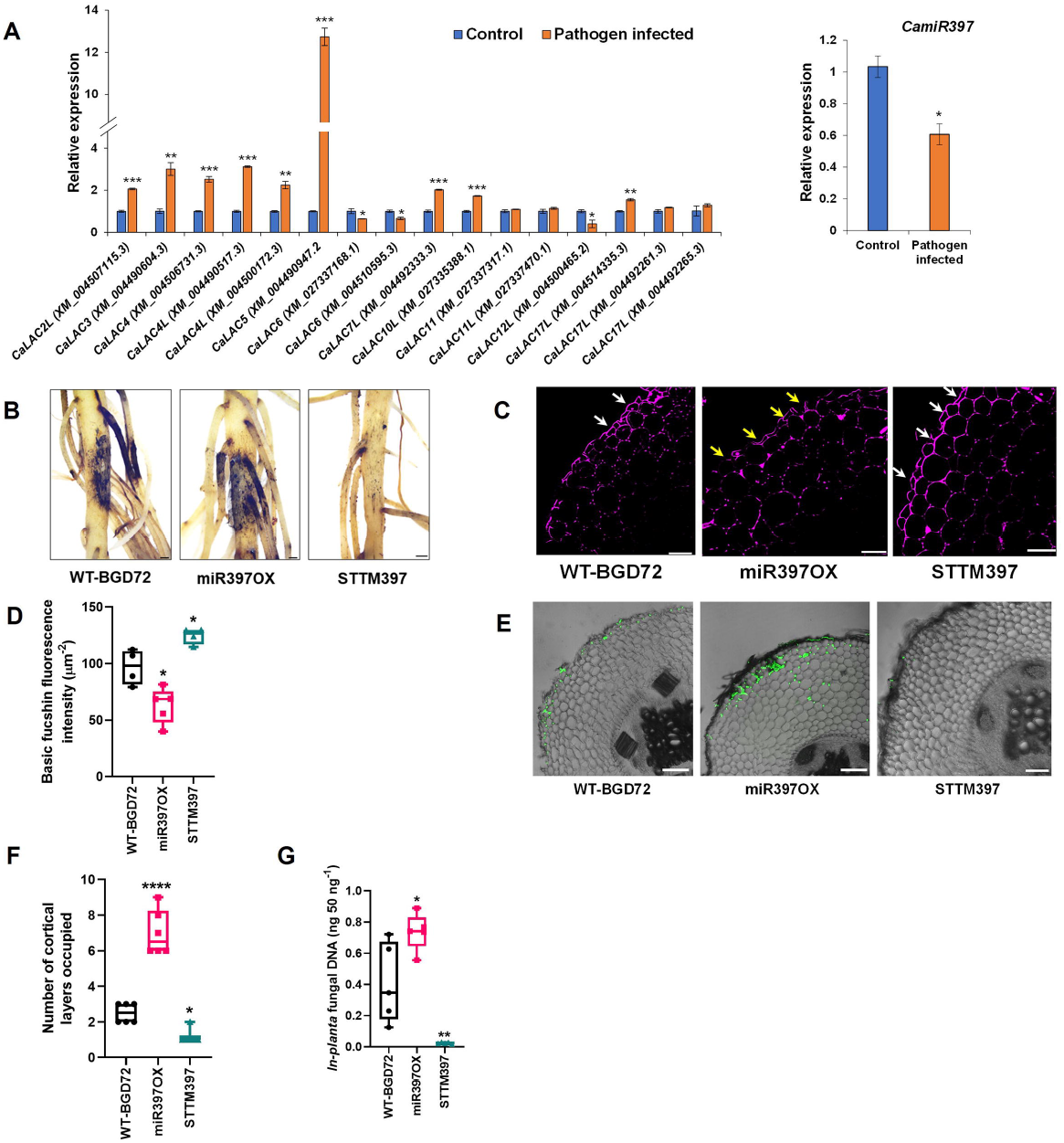
High lignin content resists the root fungal colonization. The blotting paper technique was used to impose the *Macrophomina phaseolina* infection in plant roots. Plants were examined for the infection progression 8 days after inoculation (DAI). **A**. Root specific expression of chickpea *LACCASEs* and CamiR397 in the pathogen infected BGD72 plants compared to control plants. Asterisks indicate significant differences from the control as determined by Student’s *t*-test (* for *p* < .05; ** for *p* < .01; *** for *p* < .001) **B**. Images representation to shows the presence and absence of fungal infection lesions on the roots of WT-BGD72, miR397OX (A1) and STTM397 (B4) plants. **C**. Image panel exhibiting the Transverse sections (TS) of root stained with basic fuchsin for lignin in the infected WT, miR397OX, and STTM397 plants. **D**. Graph representing the fluorescent intensity of basic fuchsin stained lignin in roots of WT, miR397OX, and STTM397 plants calculated with ImageJ software. **E**. TS of chickpea root of WT and transgenic plants exhibiting fungal infection stained with WGA-FITC. **F**. Graphical demonstration of the number of cortical cell layers colonized by fungus in WT-BGD72, miR397OX, and STTM397 plants. **G**. *In-planta* fungal DNA (ng 50 ng^-1^ total genomic DNA) estimation in the roots of infected WT-BGD72, miR397OX, and STTM397 plants. Asterisks indicate significant differences from the WT-BGD72 as determined by Student’s *t*-test (* for *p* < .05; ** for *p* < .01; *** for *p* < .001; **** for p<0.0001). Error bars show ± SE. WGA-FITC was used to label the fungal mycelia present in the roots. DAI-days after inoculation, WGA-FITC: wheat germ agglutinin-fluorescein isothiocyanate.

## Discussion

Plants are increasingly being subjected to a variety of stresses due to continuous climate changes. Accordingly, they have evolved with adaptive capabilities to alter their physiology and biochemistry (Fini *et al*., 2017). Lignin deposition in the tracheary elements of protoxylems and casparian strips in the endodermis is one such mechanism which provides mechanical strength to the cells to combat abiotic stresses (Naseer *et al*., 2012; Lee *et al*., 2013). In addition, local lignin deposition in the infected cells was also reported (Yang *et al*., 2018). Lignin deposition was also reported in the cortex’s epidermis, exodermis, and sclerenchyma cells (Peterson and Perumalla, 1990; Begovic *et al*., 2015). Besides providing mechanical strength, lignin also acts as a hydrophobic barrier to water and nutrients (Schuetz *et al*., 2014; Liu *et al*., 2017). PEROXIDASEs and LACCASEs are the two major family of enzymes involved in the final step of lignin biosynthesis. Here we report how miR397 regulates *LAC4* and *LAC17L* to increase lignin deposition in the root of a drought-tolerant chickpea variety BGD72 during natural drought and *M. phaseolina* infection.

MicroRNA397 has been studied mostly in *Arabidopsis*, poplar and rice. Ptr-miR397a overexpression in poplar downregulated abundance of 17 out of 49 *LACCASEs* and decreased lignin deposition. Out of these 17 laccases, five are homologs of AtLAC4 and six are homologs of AtLAC17 (Lu *et al*., 2013). *Arabidopsis* mutant lines lacking *LAC4* and *LAC17* expression showed 20-40 % lower lignin content in stem, but grew normally (Berthet *et al*., 2011) and *lac4lac11lac17* triple mutant lacked lignin in root (Zhao et al., 2013). Overexpression of *Setaria viridis* miR397 in *Arabidopsis* decreased *LAC4* and *LAC17* expression and lignin in stem depending on the expression level of the SvmiR397 and the overexpressing plants showed salt tolerance (Nguyen *et al*., 2020). Overexpression of *Arabidopsis* miR397b also decreased *LAC4* expression and stem lignin deposition in *Arabidopsis* (Wang *et al*., 2014). All these results clearly indicate that *LAC4* and *LAC17* are the major targets of miR397 and higher expression of this microRNA decreases expression of *LAC4* and *LAC17*, thereby, reduces lignin deposition in stem. However, role of miR397 in regulating its targets and lignin deposition in root has not been studied in crops.

Physiological response of plants towards natural drought is different from that towards the treatment with polyethylene glycol (PEG) or mannitol. Natural drought in soil is a slow process and creates a gradient of soil moisture. While treatment with PEG or mannitol immediately stops root growth in *Arabidopsis* (Khandal *et al*., 2020), drought allows cell elongation in the root elongation zone (Monteith JL, 1986; Watt *et al*., 2005; Bengough *et al*., 2011) and we observed an increase in the chickpea root length under drought. We observed a high lignin deposition in the vascular tissue within the differentiated root zone. Mannitol treatment increased root lignin deposition and also miR397b expression in *Arabidopsis* (Khandal *et al*., 2020). In contrast, we observed in the small RNA expression profile a decreased expression of CamiR397 in chickpea under natural drought. Our treatment increased expression of only ten *LACCASE*s in root probably, either those are not responsive to our treatment or they express in other tissues. Out of the three predicted CamiR397-target genes, we did not consider *LAC4L* (XM_004500172.3) for further experiment because, it did not show any significant change in response to drought and transient expression of CamiR397. We kept the names of the chickpea *LACCASE* genes following their names mentioned in the NCBI. A number of chickpea *LACCASE* genes clustered with a number of *Arabidopsis* and *Medicago LACCASEs*, and a number of chickpea (and Medicago) LACCASES having the same name create confusions. We believe the chickpea and Medicago LACCASES need new unique nomenclature, however, it is beyond the scope of this study.

Expression of Promoter-GUS constructs followed by staining showed CamiR397 and its targets are expressed mostly in the vascular tissues within the root, however, a lower expression in the cortex region was also visible. GUS staining was not visible in the cortex in the transverse root sections probably due to weak activity of the promoters in this tissue and thin sections. LAC4 and LAC17L seem to be the major enzymes among other LACCASEs involved in lignin biosynthesis in root as we see downregulation of *LAC4* and *LAC17L* led to three-fold downregulation in root lignin content. It is substantiated by the STTM397 expression which caused 1.5-fold increase in root lignin and hence, indicated that CamiR397-mediated regulation of *LAC4* and *LAC17L* is the *in-planta* mechanism of modulation of lignin synthesis and deposition in chickpea root. We observed a constitutive lignin deposition in the root exodermis. The extent of lignin deposition in exodermis followed the trend of the same in the vascular tissue in the miR397OX and STTM397 lines. MicroRNAs are labile molecules and move through tissues (Martin-Gonzalez and Suarez-Lopez, 2012). Therefore, although the expression of CamiR397 was primarily observed in the root vascular tissue and cortex, it is quite possible that it moves to other tissues to regulate LAC expression.

We have shown that lower and higher lignin deposition increased and decreased thickness of the xylem cell walls which would alter the diameter of the xylems and subsequently affect water movement. We did not observe much significant physiological ramification of the altered xylem diameter by measuring photosynthetic and biochemical parameters in the control condition except leaf relative water content. We did not observe any significant phenotypic difference between the non-transformed and the transformed plants. However, under the natural drought treatment, miR397OX plants exhibited more sensitivity than the non-transformed plants except stomatal conductance. Similarly, STTM397 plants showed more tolerance in all the parameters except in leaf relative water content.

Root is subject to various soil-borne pathogen attacks, which badly affect the crop yield. Dry root rot causes up to 70 % yield loss in chickpea under drought condition. There are several reports showing lignin deposition prevents the spread of the pathogen and the pathogen-induced lignin deposition in apple root (Hammerschmidt R., 1984; Lee *et al*., 2019; Ride, 1975). The root of the apple genotype that was resistant to *Pythium ultimum* produced more lignin upon infection with the pathogen than the sensitive genotype (Zhu and Zhou, 2021). We observed similar lignin deposition in chickpea root upon infection with the dry root rot causing pathogen *M. phaseolina*. Lower lignin deposition and subsequent spread of the pathogen in the miR397OX root suggested that CamiR397 also functions beyond the endodermis due to its expression in that tissue or due to its inter-tissue movement (Liu and Chen, 2018). It also suggested that *LACCASES* which are the targets of CamiR397 are the major *LACCASES* that forms lignin in chickpea root early in the infection by *M. phaseolina*. Our results linked miR397 with root disease resistance and showed a possible approach for introducing resistance against dry root rot in chickpea. Altogether, our results described the role of CamiR397 and its target *LACCASES* in the root lignin deposition and its physiological significance in drought response in chickpea. Additionally, our result also shed a light on the possible remedy of dry root rot in chickpea by modulating pathogen-induced lignin deposition by CamiR397 or by analysing CamiR397 expression and measuring pathogen-induced lignin deposition in different chickpea genotypes.

## Acknowledgements

The assistance of central instrumentation facilities, metabolomics facility and DISC of NIPGR in various experiments is acknowledged. The authors are grateful to the DBT-eLibrary Consortium (DeLCON) for providing access to e-Resources. The project is funded by the National Institute of Plant Genome Research, Department of Biotechnology (DBT), Ministry of Science and Technology, Government of India. DC acknowledges J.C. Bose Fellowship (JCB/2020/000014) from Science and Engineering Research Board, Department of Science and Technology. NKS and VI acknowledge DBT, SY and AF acknowledge fellowships from Council of Scientific and Industrial Research (CSIR, Govt. of India). The funders had no role in study design, data collection and analysis, decision to publish, or preparation of the manuscript. The authors have no conflicts of interest to declare.

## Authors contributions

DC initiated, conceived, designed, and coordinated the research project and wrote the manuscript. NKS designed stress related experiments, performed and analysed all the data. NKS and SKG contributed in generating chickpea transgenic lines. NKS and VI has contributed in phenotypic evaluation. VI and SY has contributed in dry root rot study. AF has done bioinformatic analysis. MSK guided the pathogen infection experiment. All authors have read and approved the final manuscript.

## Supporting Information

**Supplementary Figure S1**. Relative soil moisture content, lignin estimation under drought and RLM-RACE

**Supplementary Figure S2**. STTM construct preparation.

**Supplementary Figure S3**. Relative expression of chickpea LACCASEs in WT and transgenic lines.

**Supplementary Figure S4**. Comparison of phenotypes and agronomic parameters of the mature control and transgenic plants (T3).

**Supplementary Figure S5**. Lignin deposition in the chickpea root exodermis.

**Supplementary Figure S6**. Fungal colonization and lignin deposition in control (mock) and infected roots.

**Supplementary Table S1**. List of Chickpea Laccase genes and proteins with NCBI accession numbers

**Supplementary Table S2**. List of predicted targets of CamiR397

**Supplementary Table S3**. List of primers used in the study

**Supplementary Text S1**. Promoter sequences of CamiR397, *CaLAC4* and *CaLAC17L*

## Notes

### Competing Interest Statement

The authors have declared no competing interest.

## References

Abdel-Ghany SE, Pilon M. 2008. MicroRNA-mediated systemic down-regulation of copper protein expression in response to low copper availability in Arabidopsis. Journal of Biological Chemistry 283, 15932–15945.

Addington RN, Donovan LA, Mitchell RJ, Vose JM, Pecot SD, Jack SB, Hacke UG, Sperry JS, Oren R. 2006. Adjustments in hydraulic architecture of Pinus palustris maintain similar stomatal conductance in xeric and mesic habitats. Plant, Cell and Environment 29, 535–545.

Assfaw D, Negash T. 2020. Spatial Distribution and Association of Chickpea Wilt / Root Rots Epidemics with Biophysical Factors at West Shewa, Oromia Regional State, Ethiopia Plant Pathology & Microbiology. 11, 7–12.

Bakhshi B, Mohseni Fard E, Nikpay N, Ebrahimi MA, Bihamta MR, Mardi M, Salekdeh GH. 2016. MicroRNA signatures of drought signaling in rice root. PLoS ONE 11, e0156814.

Begović L, Ravlić J, Lepeduš H, Leljak-Levanić D, Cesar V. 2015. The pattern of lignin deposition in the cell walls of internodes during barley (Hordeum vulgare L.) development. Acta biologica cracoviensia series botanica 52, 55–66.

Bengough AG, McKenzie BM, Hallett PD, Valentine TA. 2011. Root elongation, water stress, and mechanical impedance: a review of limiting stresses and beneficial root tip traits. Journal of Experimental Botany 62, 59–68.

Berthet S, Demont-Caulet N, Pollet B, Bidzinsky P, Cezard L, Bris PL, Borrega N, Herve J, Blondet E, Balzergue Lapierre C, Jouanin L. 2011. Disruption of LACCASE4 and 17 results in tissue-specific alterations to lignification of Arabidopsis thaliana stems. Plant Cell 23, 1124–1137.

Boerjan W, Ralph J, Baucher M. 2003. Lignin biosynthesis. Annual Review of Plant Biology 54, 519–546.

Berthet S, Demont-Caulet N, Pollet B 2009. Xylem hydraulic physiology: The functional backbone of terrestrial plant productivity. Plant Science 177, 245–251.

Cai C, Xu CJ, Li X, Ferguson I, Chen KS. 2006. Accumulation of lignin in relation to change in activities of lignification enzymes in loquat fruit flesh after harvest. Postharvest Biology and Technology 40, 163–169.

Caño-Delgado A, Penfield S, Smith C, Catley M, Bevan M. 2003. Reduced cellulose synthesis invokes lignification and defense responses in Arabidopsis thaliana. Plant Journal 34, 351–362.

Berthet S, Demont-Caulet N, Pollet B 2006. Isolation and characterisation of a family of laccases in maize. Plant Science 171, 217–225.

Cárdenas W, Dankert JR. 2000. Cresolase, catecholase and laccase activities in haemocytes of the red swamp crayfish. Fish and Shellfish Immunology 10, 33–46.

Cesarino I, Araújo P, Sampaio Mayer JL, Vicentini R, Berthet S, Demedts B, Vanholme B, Boerjan W, Mazzafera P. 2013. Expression of SofLAC, a new laccase in sugarcane, restores lignin content but not S:G ratio of Arabidopsis lac17 mutant. Journal of Experimental Botany 64, 1769–1781.

Chen H, Li Y, Ma X, Guo L, He Y, Ren Z, Kuang Z, Zhang X, Zhang Z. 2019. Analysis of potential strategies for cadmium stress tolerance revealed by transcriptome analysis of upland cotton. Scientific Reports 2019 9: 86.

Chen C, Ridzon DA, Broomer AJ, Zhou Z, Lee DH, Nguyen JT, Barbisin M, Xu NL, Mahuvarkar VR, Anderson MR, Lao KQ, Lival KJ, Guegler KJ. 2005. Real-time quantification of microRNAs by stem-loop RT-PCR. Nucleic acids research 33, e179.

Cho HY, Lee C, Hwang SG, Park YC, Lim HL, Jang CS. 2014. Overexpression of the OsChI1 gene, encoding a putative laccase precursor, increases tolerance to drought and salinity stress in transgenic Arabidopsis. Gene 552, 98–105.

Diamantidis G, Effosse A, Potier P, Bally R. 2000. Purification and characterization of the first bacterial laccase in the rhizospheric bacterium Azospirillum lipoferum. Soil Biology and Biochemistry 32, 919–927.

Eggert C, Temp U, Eriksson KEL. 1997. Laccase is essential for lignin degradation by the white-rot fungus Pycnoporus cinnabarinus. FEBS Letters 407, 89–92.

Esterbauer H, Schwarzl E, Hayn M. 1977. A rapid assay for catechol oxidase and laccase using 2-nitro-5-thiobenzoic acid. Analytical Biochemistry 77, 486–494.

Fan L, Linker R, Gepstein S, Tanimoto E, Yamamoto R, Neumann PM. 2006. Progressive inhibition by water deficit of cell wall extensibility and growth along the elongation zone of maize roots is related to increased lignin metabolism and progressive stelar accumulation of wall phenolics. Plant physiology 140, 603–612.

Fini A, Tattini M, Esteban R. 2017. Editorial: Plants’ responses to novel environmental pressures. Frontiers in Plant Science 8, 2000.

Fischer KS, Johnson EC EG. 1983. Breeding and Selection for Drought Resistance in Tropical Maize. Mexico City (Mexico): CIMMYT.

Fukuda H. 2000. Programmed cell death of tracheary elements as a paradigm in plants. Plant Molecular Biology 44, 245–253.

Garg R, Patel RK, Tyagi AK, Jain M. 2011. De Novo Assembly of Chickpea Transcriptome Using Short Reads for Gene Discovery and Marker Identification. DNA Research 18, 53–63.

Garg R, Sahoo A, Tyagi AK, Jain M. 2010. Validation of internal control genes for quantitative gene expression studies in chickpea (Cicer arietinum L.). Biochemical and biophysical research communications 396, 283–288.

Gibson LJ. 2012. The hierarchical structure and mechanics of plant materials. Journal of The Royal Society Interface 9, 2749–2766.

Givaudan A, Effosse A, Faure D, Potier P, Bouillant M-L, Bally R. 1993. Polyphenol oxidase in Azospirillum lipoferum isolated from rice rhizosphere: Evidence for laccase activity in non-motile strains of Azospirillum lipoferum. FEMS Microbiology Letters 108, 205–210.

Guo Z, Kuang Z, Wang Y, Zhao Y, Tao Y, Cheng C, Yang J, Lu X, Hao C, Wang T, Cao X, Wei J, Li L, Yang X. 2020. PmiREN: a comprehensive encyclopedia of plant miRNAs. Nucleic Acids Research 48, D1114–D1121.

Hammerschmidt R. 1984. Rapid deposition of lignin in potato tuber tissue as a response to fungi non-pathogenic on potato. Physiological Plant Pathology 24, 33–42.

He F, MacHemer-Noonan K, Golfier P, Unda F, Dechert J, Zhang W, Hoffman N, Samuels L, Mansfield SD, Rausch T, Wolf S. 2019. The in vivo impact of MsLAC1, a Miscanthus laccase isoform, on lignification and lignin composition contrasts with its in vitro substrate preference. BMC Plant Biology 19, 1–18.

Hu Q, Min L, Yang X, Jin S, Zhang L, Li Y, Ma Y, Qi X, Li D, Liu H, Lindsey K, Zhu L, Zhang X. 2018. Laccase GhLac1 Modulates Broad-Spectrum Biotic Stress Tolerance via Manipulating Phenylpropanoid Pathway and Jasmonic Acid Synthesis. Plant physiology 176, 1808–1823.

Hüttermann A, Mai C, Kharazipour A. 2001. Modification of lignin for the production of new compounded materials. Applied Microbiology and Biotechnology 55, 387–394.

Irulappan V, Kandpal M, Saini K, Rai A, Ranjan A, Sinharoy S, Senthil-Kumar M. 2022. Drought Stress Exacerbates Fungal Colonization and Endodermal Invasion and Dampens Defense Responses to Increase Dry Root Rot in Chickpea. Molecular Plant Microbe Interaction 35, 583–591.

Irulappan V, Mali KV, Patil BS, Manjunatha H, Muhammad S, Senthil-Kumar M. 2021. A sick plot–based protocol for dry root rot disease assessment in field-grown chickpea plants. Applications in Plant Sciences 9, e11445.

Irulappan V, Senthil-Kumar M. 2021. Dry Root Rot Disease Assays in Chickpea: a Detailed Methodology. JoVE Journal of Visualized Experiments 167, e61702.

Jain M, Chevala VVSN, Garg R. 2014. Genome-wide discovery and differential regulation of conserved and novel microRNAs in chickpea via deep sequencing. Journal of Experimental Botany 65, 5945–5958.

Jain M, Misra G, Patel RK, Priya P, Jhanwar S, Khan AW, Shah N, Singh V, Garg R, Jeena G, Yadav M, Kant C, Sharma P, Yadav G, Bhatia S, Tyagi AK, Chattopadhyay D. 2013. A draft genome sequence of the pulse crop chickpea (Cicer arietinum L.). The Plant Journal 74, 715–729.

Jbir N, Chaïbi W, Ammar S, Jemmali A, Ayadi A. 2001. Root growth and lignification of two wheat species differing in their sensitivity to NaCl, in response to salt stress. Comptes Rendus de l’Academie des Sciences - Serie III 324, 863–868.

Jefferson RA, Kavanagh TA, Bevan MW. 1987. GUS fusions: beta-glucuronidase as a sensitive and versatile gene fusion marker in higher plants. The EMBO Journal 6, 3901.

Jeong DH, Park S, Zhai J, Gurazada SGR, de Paoli E, Meyers BC, Green PJ. 2011a. Massive analysis of rice small RNAs: Mechanistic implications of regulated MicroRNAs and variants for differential target RNA cleavage. Plant Cell 23, 4185–4207.

Khaledian Y, Maali-Amiri R, Talei A. 2015. Phenylpropanoid and antioxidant changes in chickpea plants during cold stress. Russian Journal of Plant Physiology 2015 62:6 62, 772–778.

Khandal H, Singh AP, Chattopadhyay D. 2020. The microRNA397b-LaCCASE2 module regulates root lignification under water and phosphate deficiency. Plant Physiology 182, 1387–1403.

Kim HK, Park J, Hwang I. 2014. Investigating water transport through the xylem network in vascular plants. Journal of Experimental Botany 65, 1895–1904.

Lalonde S, Wipf D, Frommer WB. 2004. Transport mechanisms for organic forms of carbon and nitrogen between source and sink. Annual Review of Plant Biology 55, 341–372.

Lee M-H, Jeon HS, Kim SH, Chung JH, Roppolo D, Lee H-J, Cho HJ, Tobimatsu Y, Ralph J, Park OK. 2019. Lignin-based barrier restricts pathogens to the infection site and confers resistance in plants. The EMBO Journal 38, e101948.

Lee Y, Rubio MC, Alassimone J, Geldner N. 2013. A mechanism for localized lignin deposition in the endodermis. Cell 153, 402–412.

Liang M, Davis E, Gardner D, Cai X, Wu Y. 2006a. Involvement of AtLAC15 in lignin synthesis in seeds and in root elongation of Arabidopsis. Planta 224, 1185–1196.

Liang M, Haroldsen V, Cai X, Wu Y. 2006b. Expression of a putative laccase gene, ZmLAC1, in maize primary roots under stress. Plant, Cell and Environment 29, 746–753.

Liu Q, Luo L, Zheng L. 2018. Lignins: Biosynthesis and biological functions in plants. International Journal of Molecular Sciences 19.

Livak KJ, Schmittgen TD. 2001. Analysis of relative gene expression data using real-time quantitative PCR and the 2(-Delta Delta C(T)) Method. Methods (San Diego, Calif.) 25, 402–408.

Lu S, Li Q, Wei H, Chang MJ, Tunlaya-Anukit S, Kim H, Liu J, Song J, Sun YH, Yuan L, Yeh TF, Peszlen I, Ralph J, Sederoff RR, Chiang VL. 2013. Ptr-miR397a is a negative regulator of laccase genes affecting lignin content in Populus trichocarpa. Proceedings of the National Academy of Sciences of the United States of America 110, 10848–10853.

Martin-Gonzalez E and Suarez-Lopez P. 2012. “And yet it moves”: Cell-to cell and long distance signalling by plant microRNAs. Plant Science 196, 18–30.

Maseda PH, Fernández RJ. 2006. Stay wet or else: Three ways in which plants can adjust hydraulically to their environment. Journal of Experimental Botany 57, 3963–3977.

Mencuccini M. 2003. The ecological significance of long-distance water transport: Short-term regulation, long-term acclimation and the hydraulic costs of stature across plant life forms. Plant, Cell and Environment 26, 163–182.

Monteith JL. 1986. How do crops manipulate water supply and demand? Philosophical Transactions - Royal Society of London, Series A 316, 245–259.

Moreira-Vilar FC, Siqueira-Soares R de C, Finger-Teixeira A, Oliveira DM de, Ferro AP, da Rocha GJ, Ferrarese M de LL, dos Santos WD, Ferrarese-Filho O. 2014. The Acetyl Bromide Method Is Faster, Simpler and Presents Best Recovery of Lignin in Different Herbaceous Tissues than Klason and Thioglycolic Acid Methods (M Reigosa, Ed.). PLoS ONE 9, e110000.

Naseer S, Lee Y, Lapierre C, Franke R, Nawrath C, Geldner N. 2012. Casparian strip diffusion barrier in Arabidopsis is made of a lignin polymer without suberin. Proceedings of the National Academy of Sciences of the United States of America 109, 10101–10106.

Nene YL, Sheila K, Sharma SB. 1996. a World List of Chickpea and Pigeonpea Pathogens. Science 59, 184–185.

Nguyen DQ, Brown CW, Pegler JL, Eamens AL, Grof CPL. 2020. Molecular Manipulation of MicroRNA397 Abundance Influences the Development and Salt Stress Response of Arabidopsis thaliana. International Journal of Molecular Sciences 21, 1–24.

Pareek A, Rathi D, Mishra D, Chakraborty S, Chakraborty N. 2019. Physiological plasticity to high temperature stress in chickpea: Adaptive responses and variable tolerance. Plant science: an international journal of experimental plant biology 289, 110258

Peterson CA and Perumalla CJ. 1990. A survey of angiosperm species to detect hypodermal casparian bands. II. Roots with a multiseriate hypodermis or epidermis. Botanical Journal of the Linnean Society. 103 (2): 113–125

Rai A, Irulappan V, Senthil-Kumar M. 2022. Dry Root Rot of Chickpea: A Disease Favored by Drought. Plant Disease 106, 346–356.

Ralph J, Lundquist K, Brunow G, Lu F, Kim H, Schatz PF, Maitra JM, Hartfield RD, Ralph SA, Christiensen JH, Boerjan W. 2004. Lignins: Natural polymers from oxidative coupling of 4-hydroxyphenyl-propanoids. Phytochemistry Reviews 3, 29–60.

Ranocha P, Chabannes M, Chamayou S, Danoun S, Jauneau A, Boudet AM, Goffner D. 2002. Laccase down-regulation causes alterations in phenolic metabolism and cell wall structure in poplar. Plant Physiology 129, 145–155.

Sachdeva S, Bharadwaj C, Singh RK, Jain PK, Patil BS, Roorkiwal M, Varshney R. 2020. Characterization of ASR gene and its role in drought tolerance in chickpea (Cicer arietinum L.). PLOS ONE 15, e0234550.

Schneider CA, Rasband WS, Eliceiri KW. 2012. NIH Image to ImageJ: 25 years of image analysis. Nature Methods 9, 671–675.

Schuetz M, Benske A, Smith RA, Watanabe Y, Tobimatsu Y, Ralph J, Demura T, Ellis B, Samuels AL. 2014. Laccases direct lignification in the discrete secondary cell wall domains of protoxylem. Plant Physiology 166, 798–807.

Sharma NK, Gupta SK, Dwivedi V, Chattopadhyay D. 2020. Lignin deposition in chickpea root xylem under drought. Plant Signaling & Behavior 15:6, 1754621.

Shen Y, Zhang Y, Chen J, Lin H, Zhao M, Peng H, Liu L, Yuan G, Zhang Z, Pan G. 2013. Genome expression profile analysis reveals important transcripts in maize roots responding to the stress of heavy metal Pb. Physiologia Plantarum 147, 270–282.

Srivastava S, Zheng Y, Kudapa H, Jagadeeswaran G, Hivrale V, Varshneyc RK, Sunkara R. 2015. High throughput sequencing of small RNA component of leaves and inflorescence revealed conserved and novel miRNAs as well as phasiRNA loci in chickpea. Plant Science 235, 46–57.

Sunkar R, Zhu JK. 2004. Novel and stress regulated microRNAs and other small RNAs from Arabidopsis w inside box sign. Plant Cell 16, 2001–2019.

Swetha C, Basu D, Pachamuthu K, Tirumalai V, Nair A, Prasad M, Shivaprasad P V. 2018. Major Domestication-Related Phenotypes in Indica Rice Are Due to Loss of miRNA-Mediated Laccase Silencing. The Plant Cell 30, 2649–2662.

Thomas BR, Yonekura M, Morgan TD, Czapla TH, Hopkins TL, Kramer KJ. 1989. A trypsin-solubilized laccase from pharate pupal integument of the tobacco hornworm, Manduca sexta. Insect Biochemistry 19, 611–622.

Thurston CF. 1994. The structure and function of fungal laccases. Microbiology 140, 19–26.

Tobimatsu Y, Schuetz M. 2019. Lignin polymerization: how do plants manage the chemistry so well? Current Opinion in Biotechnology 56, 75–81.

Tronchet M, BalaguÉ C, Kroj T, Jouanin L, Roby D. 2010. Cinnamyl alcohol dehydrogenases-C and D, key enzymes in lignin biosynthesis, play an essential role in disease resistance in Arabidopsis. Molecular Plant Pathology 11, 83–92.

Turlapati P V., Kim KW, Davin LB, Lewis NG. 2011. The laccase multigene family in Arabidopsis thaliana: Towards addressing the mystery of their gene function(s). Planta 233, 439–470.

Turner S, Gallois P, Brown D. 2007. Tracheary element differentiation. Annual Review of Plant Biology 58, 407–433.

Ursache R, Andersen TG, Marhavý P, Geldner N. 2018. A protocol for combining fluorescent proteins with histological stains for diverse cell wall components. The Plant journal□: for cell and molecular biology 93, 399–412.

Vanholme R, Demedts B, Morreel K, Ralph J, Boerjan W. 2010. Lignin biosynthesis and structure. Plant Physiology 153, 895–905.

Varkonyi-Gasic E, Wu R, Wood M, Walton EF, Hellens RP. 2007. Protocol: A highly sensitive RT-PCR method for detection and quantification of microRNAs. Plant Methods 3, 1–12.

Varshney RK, Song C, Saxena RK, et al. 2013. Draft genome sequence of chickpea (Cicer arietinum) provides a resource for trait improvement. Nature Biotechnology 2013 31:3 31, 240–246.

Wang CY, Zhang S, Yu Y, Luo YC, Liu Q, Ju C, Zhang YC, Qu LH, Lucas WJ, Wang X, Chen YQ. 2014. MiR397b regulates both lignin content and seed number in Arabidopsis via modulating a laccase involved in lignin biosynthesis. Plant Biotechnology Journal 12, 1132–1142.

Watt M, Kirkegaard JA, Rebetzke GJ. 2005. A wheat genotype developed for rapid leaf growth copes well with the physical and biological constraints of unploughed soil. Functional Plant Biology 32, 695–706.

Wei JZ, Tirajoh A, Effendy J, Plant AL. 2000. Characterization of salt-induced changes in gene expression in tomato (Lycopersicon esculentum) roots and the role played by abscisic acid. Plant Science 159, 135–148.

Wei T, Tang Y, Jia P, Zeng Y, Wang B, Wu P, Quan Y, Chen A, Li Y, Wu J. 2021. A Cotton Lignin Biosynthesis Gene, GhLAC4, Fine-Tuned by ghr-miR397 Modulates Plant Resistance Against Verticillium dahliae. Frontiers in Plant Science 12, 2624.

Xiong L, Wang RG, Mao G, Koczan JM. 2006. Identification of drought tolerance determinants by genetic analysis of root response to drought stress and abscisic acid. Plant Physiology 142, 1065–1074.

Yamaguchi M, Valliyodan B, Zhang J, Lenoble ME, Yu O, Rogers EE, Nguyen HT, Sharp RE. 2010. Regulation of growth response to water stress in the soybean primary root. I. Proteomic analysis reveals region-specific regulation of phenylpropanoid metabolism and control of free iron in the elongation zone. Plant, Cell & Environment 33, 223–243.

Yang C, Liang Y, Qiu D, Zeng H, Yuan J, Yang X. 2018. Lignin metabolism involves Botrytis cinerea BcGs1-induced defense response in tomato. BMC Plant Biology 18, 1–15.

Zhang K, Lu K, Qu C, Liang Y, Wang R, Chai Y, Li J. 2013. Gene Silencing of BnTT10 Family Genes Causes Retarded Pigmentation and Lignin Reduction in the Seed Coat of Brassica napus. PLoS ONE 8, e61247

Zhang YC, Yu Y, Wang CY, Li ZY, Liu Q, Xu J, Liao JY, Wang XJ, Qu LH, Chen F, Xin P, Yan C, Chu J, Li HQ, Chen YQ. 2013. Overexpression of microRNA OsmiR397 improves rice yield by increasing grain size and promoting panicle branching. Nature Biotechnology 31, 848–852.

Zhao Q, Nakashima J, Chen F, Yin Y, Fu C, Yun J, Shao H, Wang X, Wang ZY, Dixon RA. 2013. LACCASE is necessary and nonredundant with PEROXIDASE for lignin polymerization during vascular development in Arabidopsis. Plant Cell 25, 3976–3987.

Zhou L, Liu Y, Liu Z, Kong D, Duan M, Luo L. 2010. Genome-wide identification and analysis of drought-responsive microRNAs in Oryza sativa. Journal of Experimental Botany 61, 4157–4168.

Zhu Y, Zhou Z. 2021. The genotype-specific laccase gene expression and lignin deposition patterns in apple root during Pythium ultimum infection. Fruit Research 2021 1:1 1, 1–9.

Zuker M. 2003. Mfold web server for nucleic acid folding and hybridization prediction. Nucleic Acids Research 31, 3406–3415.

