## Supplementary Materials for "MicroRNA397 regulates tolerance to drought and fungal infection by regulating lignin deposition in chickpea root"

### Supplementary data

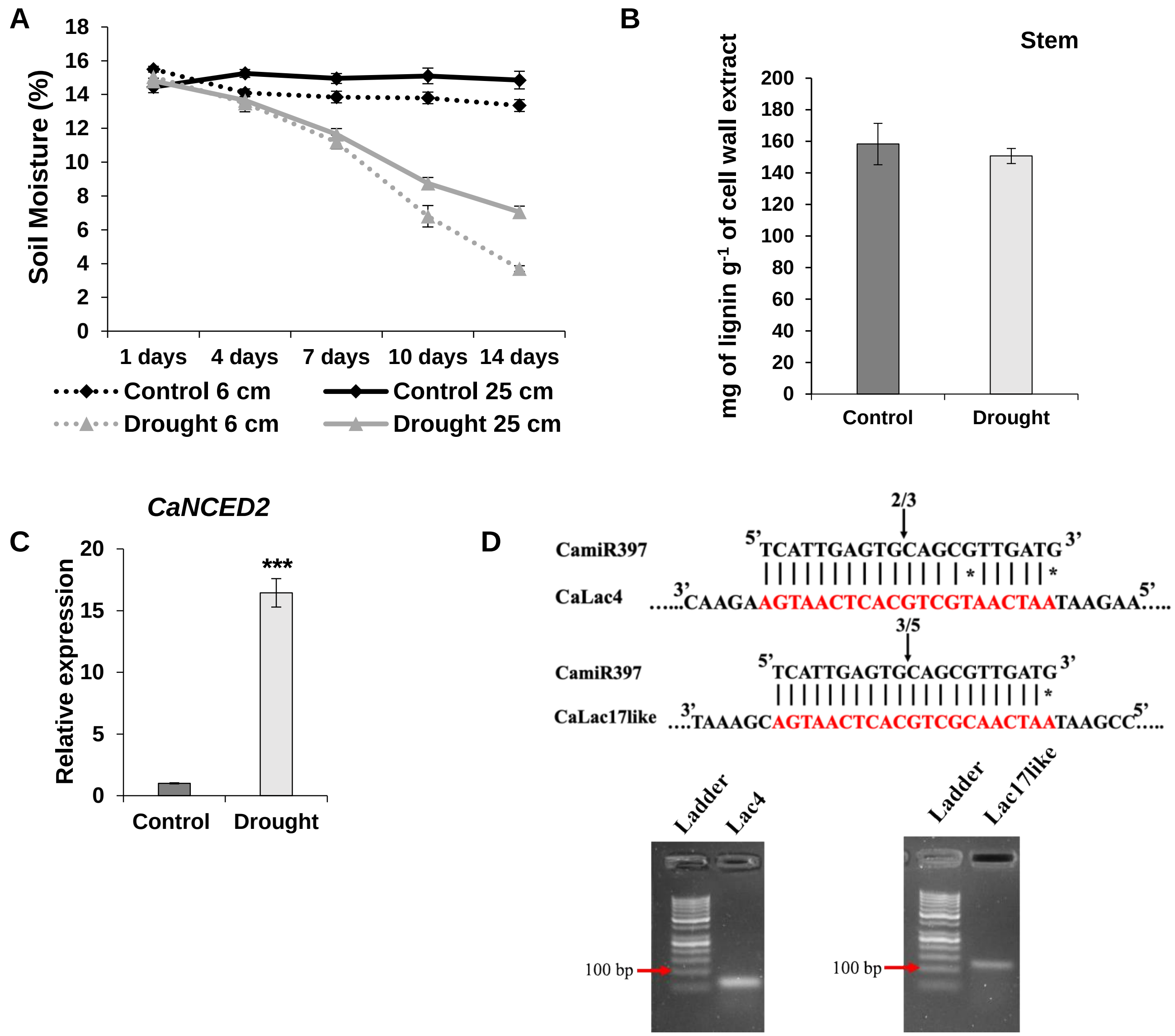


**Supplementary Figure S1. Relative soil moisture content, lignin estimation under drought and RLM-RACE A.** Relative soil moisture contents of the pot grown chickpea cultivars BGD measured from the 15th day onwards up to the 28th day upon continuation (Control) or termination (Drought) of irrigation in the drought experiment. Measurement of relative soil moisture contents were performed using soil moisture meter (Lutron PMS-714, Taiwan) at two different soil depth (6 cm and 25 cm) at three different places in the pot were sown. **B.** Lignin estimation in stem of control and drought treated plants. **C.** Relative expression level of ABA synthesis gene, 9-*cis-EPOXYCAROTENOID DIOXYGENASE* (*CaNCED2*; XM_004504855) in the root of chickpea cultivar BGD72 under control and drought condition. Plants were grown 14 days with normal hydration. Samples were harvested for analysis after 14 days of receding water condition. Asterisks indicate significant differences from the control as determined by Student's *t*-test (* for $p < .05$; ** for $p < .01$; *** for $p < .001$). Error bars show ± SE. **D.** RLM-RACE validation of cleavage of target mRNAs of CamiR397. The arrows above the sequences indicate the cleavage sites. The numbers indicate the numbers out of total molecules showed cleavage by sequencing. The images below show the amplified RLM-RACE products.

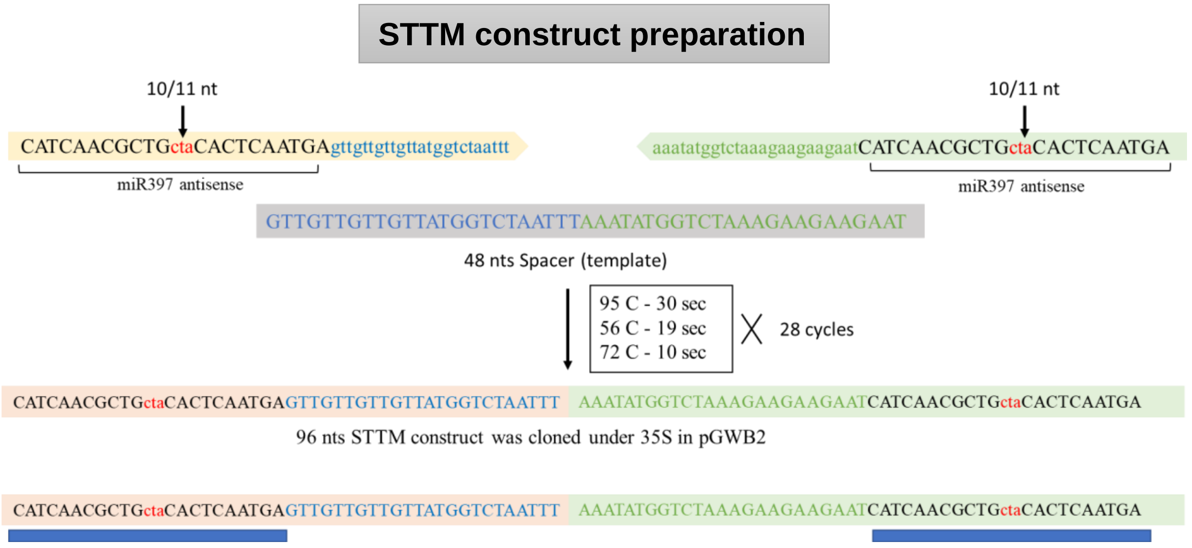


**Supplementary Figure S2. STTM construct preparation.** The target mimic construct also known as **S**hort **T**andem **T**arget **M**imic is prepared according to Yan *et al*., 2012. A 48-base pair (core) adapter with antisense CamiR397 at its ends were prepared using PCR having three nucleotides (CTA) inserted at 10/11$^{th}$ position in both antisense sequences. The CamiR397s in plants will bind to this transcript due to sequence complementarity but unable to cleave it since loss of cleavage specific site. Primer details are listed in supplementary table 3. The resultant oligonucleotide was then cloned via pENTR vector to binary vector pGWB2.

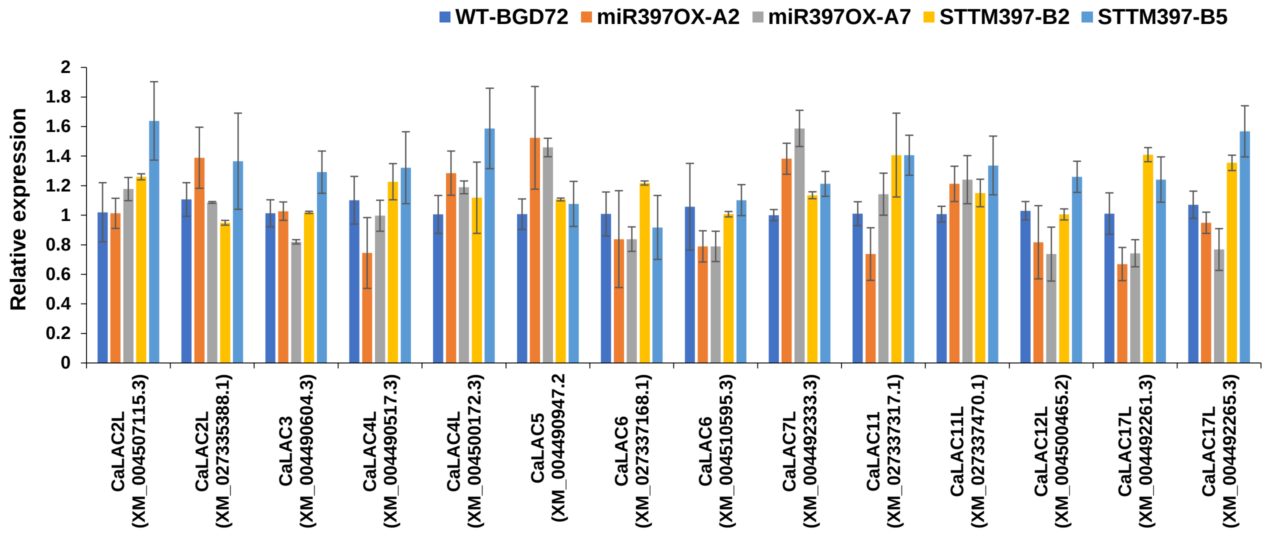


**Supplementary Figure S3. Relative expression of chickpea LACCASEs in WT and transgenic lines.** Root specific expression profile of chickpea laccases except the target laccases of CamiR397 in wild type (WT-BGD72), miR397OX (A1 & A6) and STTM397 (B2 & B4) lines through qRT-PCR. Three biological replicates were taken for this analysis. Error bars show ± SE.

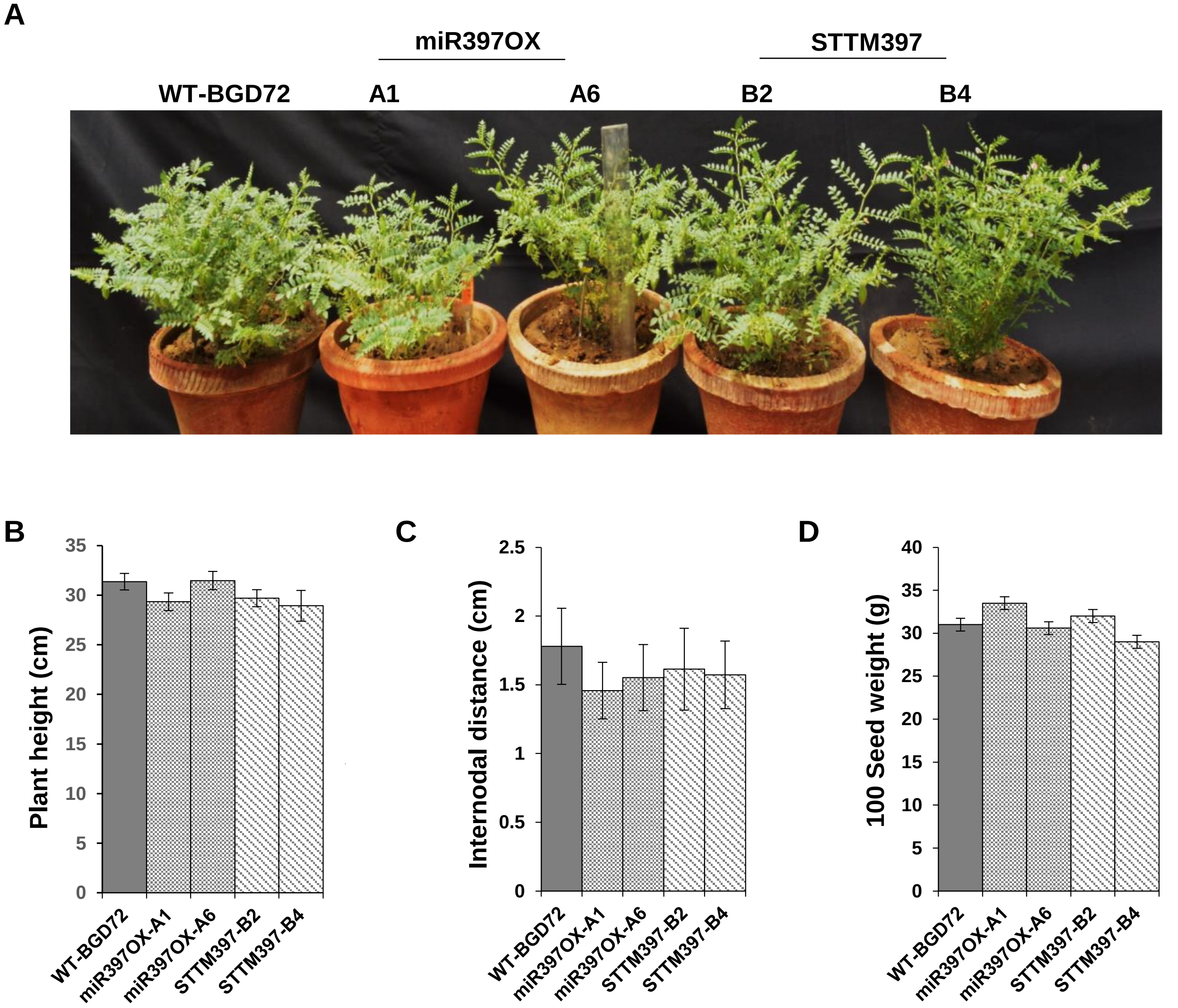


**Supplementary Figure S4. Comparison of phenotypes and agronomic parameters of the mature control and transgenic plants (T3). A.** Phenotypic observation of matured miR397OX and STTM397 transgenic chickpea plants grown under control conditions. **Agronomic parameters in chickpea transgenic lines compared to WT-BGD72**. **B.** Plant height (in cm) **C.** Internodal distance (in cm). **D.** Seed weight (in g). Plants were grown under control conditions to the full maturity. Error bars show ± SE (n=3).

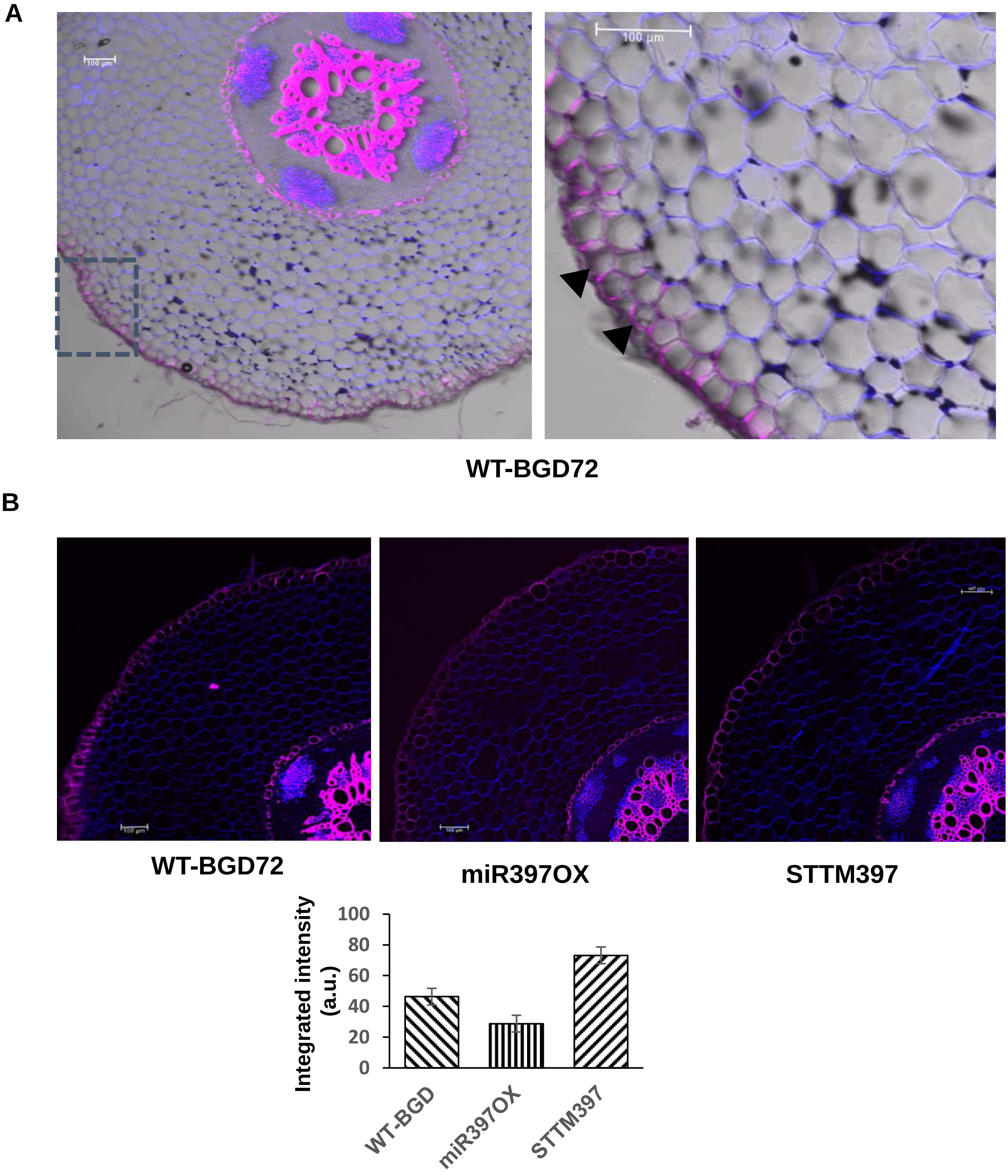


**Supplementary Figure S5. Lignin deposition in the chickpea root exodermis.** A. Transverse root section stained with basic fuchsin and sodium calcoflouor. The Flurescent images was superimposed on the bright field image. The right panel is the magnified image of the inset in the left panel. B. Transverse root sections of mentioned root samples stained with basic fuchsin and sodium calcoflouor. Scale bar 100 µm. The lower panel shows a comparison of the fluorescence intensity of basic fuchsin stain in the exodermis. Arbitrary unit was used for the comparison. The standard deviation represents data from biological replicates (n=5).

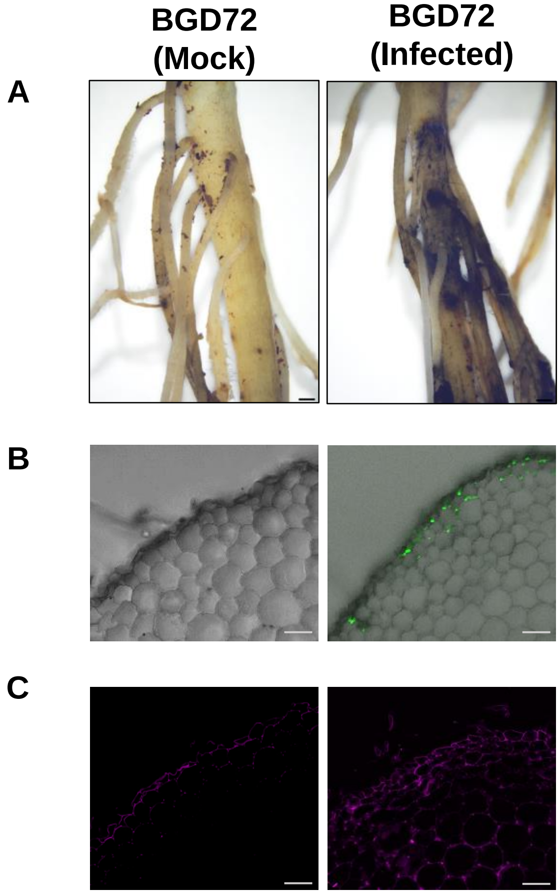


**Supplementary Figure S6. Fungal colonization and lignin deposition in control (mock) and infected roots.** Chickpea genotype (BGD72) was infected with *M. phaseolina* using the blotting paper technique. On 8 DAI, roots were collected and sections were made and stained with WGA-FITC and basic fuschin. **A.** Images representation to shows the presence and absence of fungal infection lesions on the roots of BGD72 in control and infected plants. **B.** Transverse sections (TS) of root of control and infected plants exhibiting fungal infection stained with WGA-FITC. **C.** Image panel exhibiting the TS of root stained with basic fuchsin for lignin in the both control and infected plants. Scale bar= 1 cm (A), 50µm (B & C). DAI-days after inoculation, WGA-FITC: wheat germ agglutinin-fluorescein isothiocyanate
